# Pluripotency-Independent Induction of Human Trophoblast Stem Cells from Fibroblasts

**DOI:** 10.1101/2021.11.10.468044

**Authors:** Moriyah Naama, Ahmed Radwan, Valery Zayat, Shulamit Sebban, Moran Rahamim, Rachel Lasry, Mohammad Jaber, Ofra Sabag, Hazar Yassen, Dana Orzech, Areej Khatib, Silvina Epsztejn-Litman, Michal Novoselsky-Persky, Kirill Makedonski, Noy Deri, Debra Goldman-Wohl, Howard Cedar, Simcha Yagel, Rachel Eiges, Yosef Buganim

## Abstract

Recent studies demonstrated that human trophoblast stem-like cells (hTS-like cells) can be derived from naïve embryonic stem cells or be induced from somatic cells by the pluripotency factors, OSKM. This raises two main questions; (i) whether human induced TSCs (hiTSCs) can be generated independently to pluripotent state or factors and (ii) what are the mechanisms by which hTSC state is established during reprogramming. Here, we identify GATA3, OCT4, KLF4 and MYC (GOKM) as a pluripotency-independent combination of factors that can generate stable and functional hiTSCs, from both male and female fibroblasts. By using single and double knockout (KO) fibroblasts for major pluripotency genes (i.e. SOX2 or NANOG/PRDM14) we show that GOKM not only is capable of generating hiTSCs from the KO cells, but rather that the efficiency of the process is increased. Through H3K4me2 and chromatin accessibility profiling we demonstrate that GOKM target different *loci and genes* than OSKM, and that a significant fraction of them is related to placenta and trophoblast function. Moreover, we show that GOKM exert a greater pioneer activity compared to OSKM. While GOKM target many specific hTSC *loci*, OSKM mainly target hTSC *loci* that are shared with hESCs. Finally, we reveal a gene signature of trophoblast-related genes, consisting of 172 genes which are highly expressed in blastocyst-derived TSCs and GOKM-hiTSCs but absent or mildly expressed in OSKM-hiTSCs.

Taken together, these results imply that not only is the pluripotent state, and SOX2 specifically, not required to produce functional hiTSCs, but that pluripotency-specific factors actually interfere with the acquisition of the hTSC state during reprogramming.

## INTRODUCTION

For an extensive period of time, all attempts to isolate and propagate human TSCs (hTSCs) *in vitro* had failed due to lack of knowledge of the culture conditions required for the maintenance of these cells. Recently, such culture conditions were identified and for the first time hTSCs were successfully derived and propagated from blastocysts and first trimester trophoblasts ^1^. Following differentiation, these hTSCs gave rise to all major trophoblast cell types, exhibited transcriptional and epigenetic signatures similar to first trimester cytotrophoblasts and formed trophoblast lesions when injected into NOD/SCID mice, suggesting fully functional hTSCs ^1^. However, at the present time, these cells can be isolated only during the first-trimester, while placental disorders are detected only at late stages of pregnancy. This constraint largely restricts the usefulness of these cells in modelling placental pathologies and identifying risk factors at early stages of implantation, as it is currently impossible to derive hTSCs from disease-affected term placentas ^1^.

Alternatively, the ability to convert fibroblasts into other cell types ^2^ by a defined number of transcription factors opens an attractive avenue which resolves this limitation, as mesenchymal cells can be isolated relatively easily from post-gestational tissue, such as term placenta or cord blood following disease-affected pregnancies.

Recent studies demonstrated that human induced trophoblast stem cells (hiTSCs) can be generated either by transdifferentiating human naïve pluripotent cells ^3–8^ or by forced expression of OCT4, SOX2, KLF4 and MYC (OSKM) in fibroblasts ^3, 9^. In these approaches the efficiency and quality of the cells was dependent on the initial acquisition of pluripotency and the use of naïve human embryonic stem cell (hESC) culture conditions or on the pluripotency OSKM factors. Thus, it is still unclear whether human TSC state can be directly induced from somatic cells, and which are the major factors that can mediate this process. Moreover, we and others have shown in the mouse system that the direct conversion of fibroblasts into mouse iTSCs is superior to transdifferentiation from pluripotent cells, as the latter was shown to generate unstable colonies with epigenetic abnormalities ^10–13^.

Here, we developed a new paradigm in which human fibroblasts are directly converted into human induced trophoblast stem cells (hiTSCs) by transient upregulation of GATA3, OCT4, KLF4 and MYC. hiTSCs were stable for long passages, differentiate into the various trophoblast cell types, generate trophoblastic lesions when inject into NOD/SCID mice, retain normal karyotype and form functional trophoblastic organoids. Transcriptional and methylation analyses indicate that hiTSCs closely resemble human blastocyst-derived TSCs (hbdTSCs).

Moreover, by profiling the transcriptome and chromatin (i.e. ATAC-seq and ChIP-seq for H3K4me2) at an early stage of reprogramming, we show that cells transduced with GOKM are more directed and follow a distinct route toward the TSC state compared with OSKM. Finally, we show that GOKM are more efficient in producing hiTSCs than OSKM and that the TSC state achieved by GOKM bypasses pluripotency. These data suggest that fully functional hiTSCs can be directly generated from human fibroblasts and propose a mechanism by which GOKM target the genome to acquire the hTSC state.

## RESULTS

### Ectopic expression of GATA3, OCT4, KLF4 and MYC in fibroblasts generates stable colonies that are similar in their morphology and marker expression to human trophoblast stem cells

Previously, we and others have shown that transient ectopic expression of four mouse key trophectoderm (TE) genes, *Gata3*, *Eomes*, *Tfap2c* and *Myc/Ets2* can force fibroblasts to become stable and fully functional mouse induced trophoblast stem cells (miTSCs, ^10, 13^). However, current knowledge suggests that key TE genes vary significantly between human and mouse. Single-cell RNA-seq studies of the human pre-implantation blastocyst revealed that key mouse TE genes such as *Eomes* and *Elf5* are absent or expressed at very low levels in the human TE ^14, 15^. *Esrrb*, which is expressed in the mouse epiblast ^16^ and which plays an important role in the maintenance and induction of pluripotency ^11, 17, 18^ and mTSCs ^19^, is not expressed in the human epiblast, but rather in the TE and primitive endoderm (PE) ^14, 15^. Another crucial difference between mouse and human blastocysts is the involvement of pluripotency genes such as the key master regulator OCT4 in the establishment of the human TE compartment ^20^.

Thus, in order to reprogram fibroblasts into human induced trophoblast stem cells (hiTSCs), we selected transcription factors with a known role in the development of the human trophoblast lineage but also included pluripotent genes based on their suspected necessity for the induction of the hTSC state ^20–22^. In total, we cloned seven genes, *GATA3*, *TFAP2C*, *ESRRB*, *OCT4*, *KLF4*, *SOX2* and MYC, into doxycycline (dox)-inducible lentiviral vectors and used them to infect human foreskin fibroblasts (HFFs). Cells were kept in low (5%) oxygen conditions and treated with dox for two weeks in basic reprogramming medium (DMEM + 10%FBS) which was gradually switched to hTSC medium (^1^, Figure 1A). Following 4 weeks of reprogramming, the induced cells were weaned off dox and allowed to stabilize for 7-10 days, after which individual epithelial-like colonies were manually transferred into separate plates for propagation and analysis. Transgene integration analysis revealed that *GATA3*, *OCT4*, *KLF4* and *MYC* (GOKM) were the only transgenes which had been integrated in all examined colonies (Figure S1A), suggesting that the pluripotent gene *SOX2* is not required for the induction of the TSC state.

**Figure 1.**
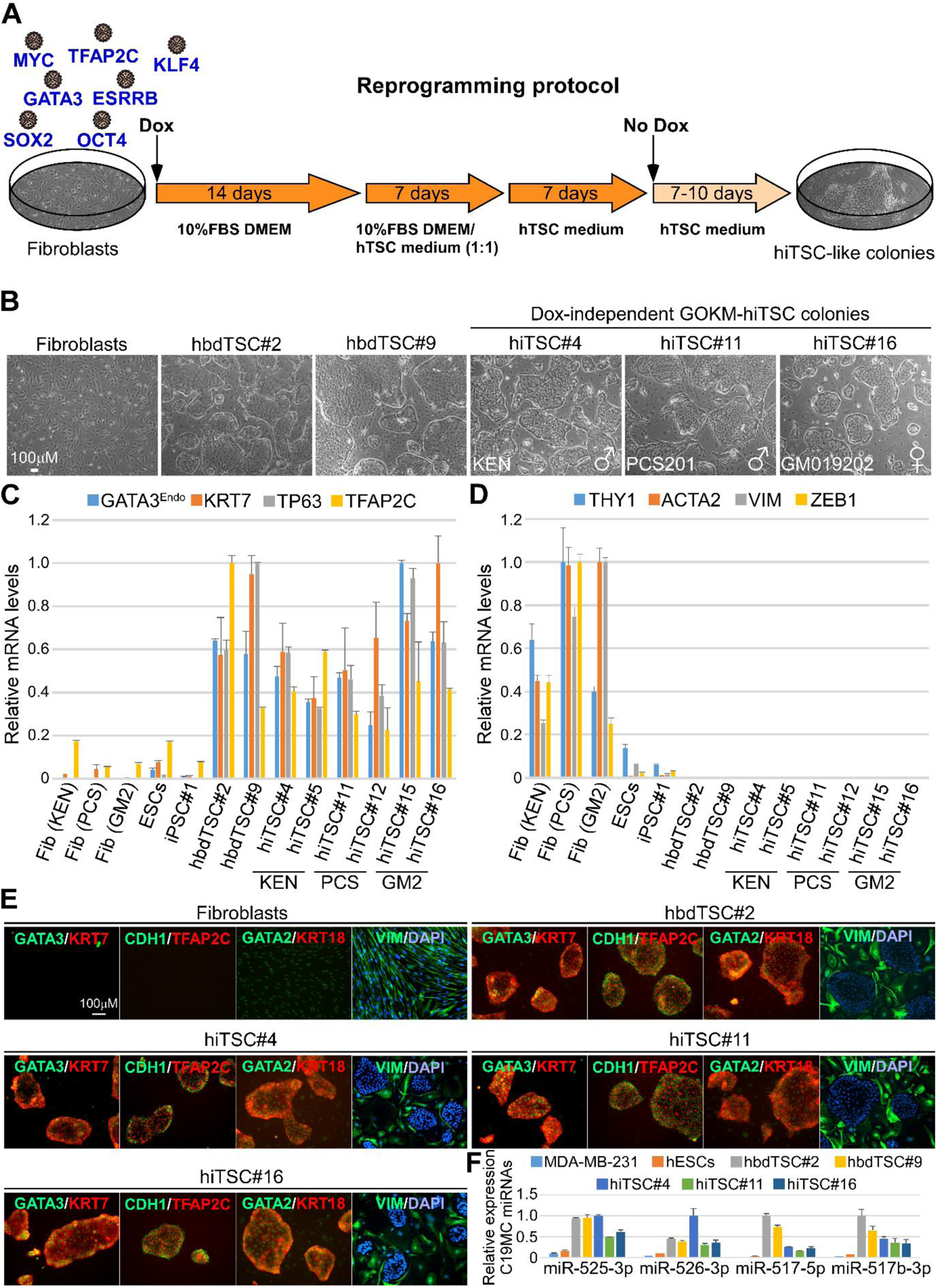
Ectopic expression of GATA3, OCT4, KLF4 and MYC (GOKM) convert human fibroblasts into trophoblast stem-like cells. **(A)** Schematic representation of the protocol for reprogramming human fibroblasts into human induced trophoblast stem cells (hiTSCs). M2rtTA-transduced fibroblasts, passage 7-14, were infected with lentiviral vectors encoding for the indicated transcription factors. Infected fibroblasts were exposed to doxycycline (dox) for 28 days, while the reprogramming medium was gradually changed as depicted in the scheme. 7-10 days post dox withdrawal, stable epithelial colonies were isolated and seeded on plates containing MEF feeder-cells. Colonies were passaged until full stabilization. **(B)** Bright field images of human primary fibroblasts, two human blastocyst-derived TSC lines, hbdTSC#2 and hbdTSC#9, and three representative GOKM-derived hiTSC colonies originating either from KEN (foreskin fibroblasts, hiTSC#4), PCS201 (foreskin fibroblasts, hiTSC#11) or GM25432 (GM2, female adult fibroblasts, hiTSC#16). **(C-D)** qPCR analysis of mRNA levels for TSC-specific markers *GATA3 (endogenous 5’ UTR expression), KRT7*, *TP63* and *TFAP2C* (C) and mesenchymal-specific markers *THY1*, *ACTA2*, *VIM* and *ZEB1* (D) in six hiTSC colonies, two hbdTSC lines, three fibroblast lines, hESCs and iPSCs. The indicated hiTSC colonies were derived from three independent reprogramming experiments. The highest sample for each gene was set to 1. Results were normalized to the mRNA levels of the housekeeping control gene *GAPDH* and are shown as fold change. Error bars indicate standard deviation between two duplicates. **(E)** Immunofluorescent staining for TSC-specific markers GATA3, KRT7, TFAP2C, GATA2, epithelial markers CDH1 and KRT18, and the mesenchymal marker VIM in parental fibroblasts (KEN) and hbdTSC#2 controls and in three representative hiTSC clones, hiTSC#4, hiTSC#11 and hiTSC#16. **(F)** qPCR analysis for the expression of *C19MC miRNA cluster* in the indicated samples. The highest sample for each *miR* was set to 1. Results were normalized to the expression levels of the control *miR 103a* and are shown as fold change. Error bars indicate standard deviation between two duplicates.

Indeed, transduction of GOKM into two primary HFF lines, namely KEN and PCS201, and one primary adult female patient-derived fibroblast line (GM2), produced stable and dox-independent (Figure S1B) epithelial-like colonies that exhibited a morphology remarkably similar to that of mTSCs and to human blastocyst-derived TSCs (hbdTSCs) following passaging on mouse feeder cells (Figure 1B). The reprogramming efficacy ranged between 2×10^−6^-5X10^−5^, depending on the origin and age of the parental fibroblasts, cell passage and infection efficiency, yielding ~5-100 colonies out of 2×10^6^ seeded cells in a 10cm plate (Figure S1C).

In order to evaluate the identity of the resultant colonies, expression of hTSC markers was assessed. Quantitative PCR (qPCR) revealed active transcription of known trophoblast markers such as *GATA2*, *TFAP2A*, *TFAP2C*, *KRT7* and *TP63*, as well as endogenous expression of *GATA3*, in a manner which is comparable to hbdTSCs (Figures 1C and S1D). As expected, the resultant hiTSC colonies showed drastic downregulation of mesenchymal markers and upregulation of epithelial markers, indicating successful mesenchymal-to-epithelial transition (MET) (Figure 1D and S1E). Of note, the epithelial marker KRT18 discriminated between human epithelial cells from trophectodermal origin (i.e. hbdTSCs and hiTSCs), in which the expression was high, and epithelial cells from a pluripotency origin (i.e. ESCs and iPSCs), similar to mouse cells ^10^.

Human trophoblast is known to have a unique expression pattern of HLA proteins, including a lack of all HLA class I molecules in villous trophoblast and lack of HLA-A expression in all known trophoblast subtypes ^23^. In accordance with that, expression of the HLA class I gene *HLA-A* was absent from all hiTSC and hbdTSC lines (Figure S1F, ^24^).

Expression of hTSC markers GATA3, GATA2, TFAP2C and KRT7, epithelial markers CDH1 and KRT18, as well as the absence of the mesenchymal marker VIM and classical HLA class I proteins (HLA-A/B/C) were validated at the protein level as well (Figures 1E and S1G). Finally, the TSC-specific C19MC miRNA cluster was highly expressed in all hiTSC colonies and hbdTSC positive controls, but not in hESCs or the breast cancer cell line MDA-MB-231 negative controls (Figure 1F). Of note, although highly expressed in all hiTSC lines, miR-517-5p demonstrated reduced levels in hiTSCs compared to hbdTSCs. Taken together, these data suggest that transient GOKM expression can force human fibroblasts to become stable, dox-independent epithelial colonies resembling hbdTSCs in their morphology and hTSC marker expression.

### The transcriptome of hiTSCs is highly similar to hbdTSCs

Extensive nuclear reprogramming during somatic cell conversion should ideally result in the activation of the newly established endogenous gene expression circuitry of the targeted cells ^25, 26^. Incomplete activation of the endogenous circuitry will lead to a partially analogous transcriptome, as can be seen in several direct conversion models ^25^. To assess whether hiTSCs successfully activated the endogenous TSC circuitry, we subjected seven hiTSC clones (hiTSC#1, hiTSC#4, hiTSC#7, hiTSC#11, hiTSC#13, hiTSC#15, hiTSC#16) to RNA-sequencing (RNA-seq) analysis. Two hbdTSC clones (hbdTSC#2 and hbdTSC#9), two parental primary fibroblast lines (KEN and GM2) and two pluripotent stem cell (PSC) clones (hESCs and hiPSC#1) were used as positive and negative controls, respectively. As OSKM factors were recently shown to be capable of producing hiTSCs as well ^3, 9^, we sought to understand whether the selection of factor combination has any effect on gene expression in the resulting hiTSCs. To that end, we reprogrammed fibroblasts into hiTSCs using the OSKM factors and profiled the transcriptome of two OSKM-hiTSC clones (OSKM-hiTSC#1, OSKM-hiTSC#2). Notably, the various hiTSC clones clustered together with hbdTSC clones and far away from the fibroblasts and hESC/hiPSC controls, as indicated by principal component analysis (PCA, Figure 2A) and hierarchical correlation heatmap (Figure 2B).

**Figure 2.**
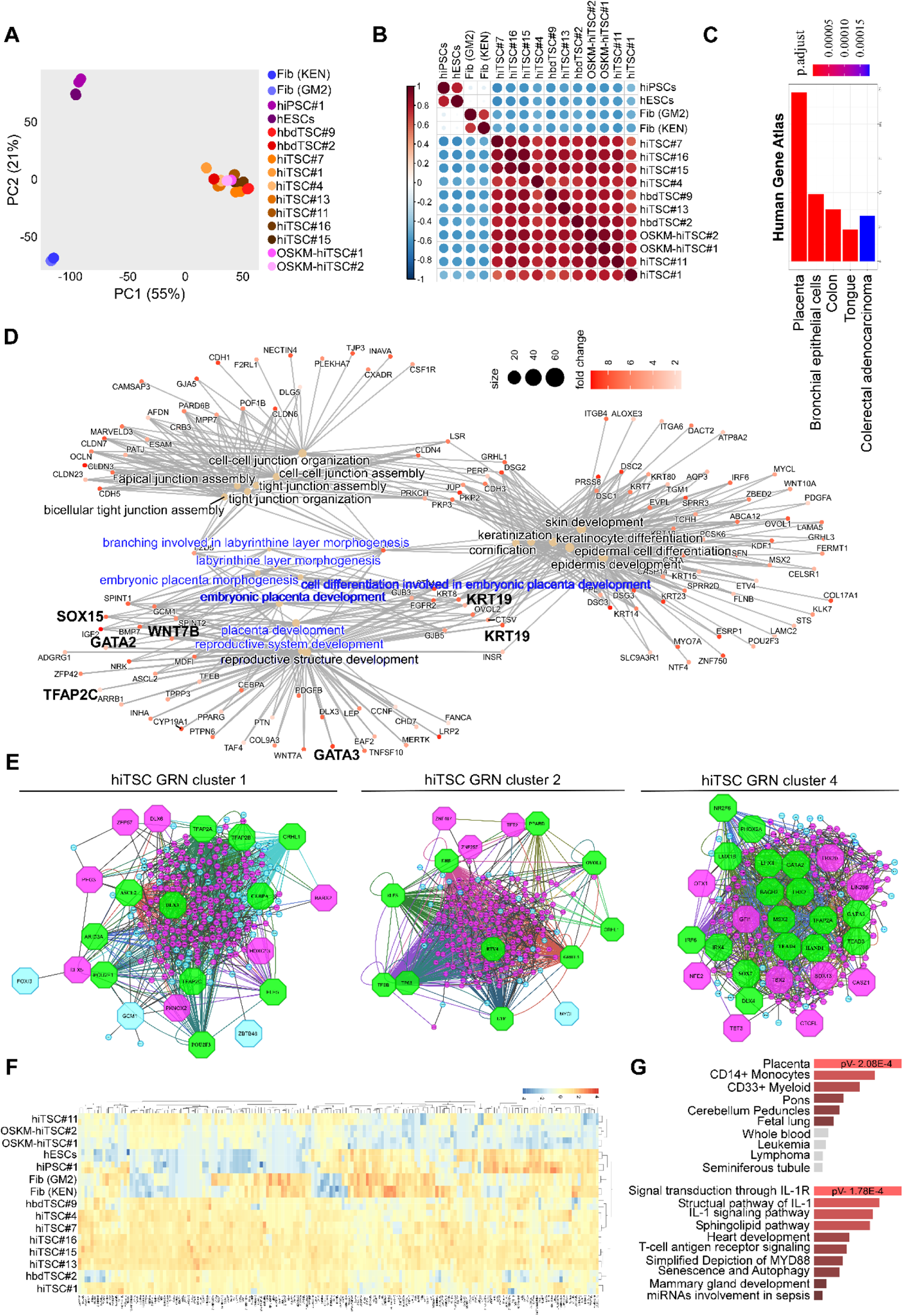
RNA-seq analysis indicates that GOKM-derived hiTSCs exhibit a transcriptome that is highly similar to that of hbdTSCs. **(A-B)** Plots based on RNA-seq data portraying comparisons of whole transcriptome of two biological duplicates of two lines of parental fibroblasts (KEN and GM2), two pluripotent stem cell clones, hESCs and hiPSCs, two hbdTSCs lines, hbdTSC#2 and hbdTSC#9, seven GOKM-derived hiTSC clones, hiTSC#1, hiTSC#4, hiTSC#7, hiTSC#11, hiTSC#13, hiTSC#15 and hiTSC#16, and two OSKM-derived hiTSC clones, OSKM-hiTSC#1 and OSKM-hiTSC#2. Principal component analysis (PCA) plot (A) constructed using the top 2000 differentially expressed genes, and correlation heatmap (B) of bulk RNA displaying the transcriptional similarity between hbdTSCs and hiTSCs and their dissimilarity from PSCs (ESCs and iPSCs) and fibroblasts (KEN and GM2). **(C)** Bar graph showing the highest enriched GO terms for the top 1000 most differentially expressed genes under the Human Gene Atlas category. **(D)** Gene concept network for top 1000 upregulated genes in hiTSCs versus fibroblasts. Genes are represented with a heat colorimetric scale showing the fold change in their expression (red for the highest fold change) and linked to up-regulated GO-terms. A significant enrichment for GO terms relevant to placenta and embryonic placenta morphogenesis and development (according to the Human Gene Atlas, marked by blue) is shown. **(E)** Network analysis for top 2000 upregulated genes in hiTSCs versus fibroblasts. Protein-protein interaction network was analyzed with STRING (http://www.string-db.org). The MCODE plugin tool in Cytoscape was used for further analysis of densely connected genes. For each subnetwork we used iRegulon plugin tool in Cytoscape to systematically analyze the composition of the gene promoters in transcription factor binding sites. **(F)** Heatmap and hierarchical clustering of 172 genes that were found to be differentially expressed in OSKM-derived hiTSCs when compared to hbdTSCs and GOKM-derived hiTSC clones. Importantly, a reciprocal analysis searching for differentially expressed genes in GOKM-hiTSCs identified only one gene (SYK) that is aberrantly expressed in GOKM-hiTSCs. **(G)** Bar graphs showing the most enriched GO terms for the 172 genes from (F), under the Human Gene Atlas and WikiPathways categories using EnrichR.

Scatter plot analysis indicated a highly similar transcriptome between hbdTSCs and hiTSCs with R^2^ scores ranging between 0.78-0.84, and key hTSC genes such as TP63 and GATA3 showing high levels of expression in all TSC samples but not in hESC or fibroblast negative controls (Figure S2A). The ~1000 most differentially expressed genes between GOKM-hiTSCs and fibroblasts revealed significant enrichment for gene ontology (GO) terms associated with placenta, trophoblast, embryonic placenta morphogenesis and development, cell-cell adhesion and response to estrogen, and identified TSC factors such as TP63, SUZ12 and GATA family genes as key transcription factors that regulate this gene list according to EnrichR database. NANOG is also featured as a key regulator, likely as a repressor. (Figures 2C-D and S2B-E). Moreover, using STRING and iRegulon for top 2000 differentially expressed genes between hiTSC and fibroblasts we identified gene regulatory networks (GRNs) and protein-protein interactions that are highly associated with the hTSC state ^22, 23, 27^. Among the key regulators of these GRNs are TP63, GATA2, GRHL3, TFAP2A, TFAP2C, ARIDA3, ELF5, TEAD4, KLF5, ETV4, ASCL2, HAND1 and many others (Figure 2E, marked by green octagons).

To test whether small differences in gene expression exist between hiTSC clones and hbdTSCs, we compared the transcriptome of OSKM-hiTSCs and GOKM-hiTSCs to hbdTSCs. While the comparison between GOKM-hiTSC clones to hbdTSCs identified only one gene (i.e. SYK) with significant differences that is aberrantly expressed in GOKM-hiTSCs, the comparison between OSKM-hiTSC clones to hbdTSCs identified 172 genes (p-value < 0.01) which were aberrantly expressed in OSKM-hiTSC clones, but not in GOKM-hiTSC clones (excluding GOKM colony 11, Figure 2F). GO term analysis revealed that this group of genes is significantly enriched for placenta and IL-1R signal transduction (Figure 2G). It is important to note that appropriate IL-1 signaling activity has been implicated in blastocyst implantation and trophoblast motility ^28, 29^. All together, these data suggest that the transcriptome of hiTSCs is highly similar to that of blastocyst-derived hTSCs, although a small difference in the expression of placenta-related genes can be seen in OSKM-hiTSCs.

### GOKM and OSKM remodel the chromatin differently at early stages of reprogramming

Given that OSKM are capable of producing hiTSCs, we next sought to understand whether GOKM and OSKM remodel the somatic chromatin in a similar manner during reprogramming. To that end, we profiled the transcriptome and chromatin of cells transduced with GOKM or OSKM using RNA-seq, Assay for Transposase-Accessible Chromatin using sequencing (ATAC-seq) and chromatin immunoprecipitation and sequencing (ChIP-seq) for the histone mark H3K4me2. We chose to profile day 3 (D3) of reprogramming because, in contrast to later stages in reprogramming, cells respond relatively homogenously to transgene induction at this time point ^26^, allowing more accurate bulk analysis. We selected the histone mark H3K4me2 because it has been shown to mark closed regions of the genome that are destined to become open later in the reprogramming process ^26, 30^. Parental fibroblasts, hbdTSCs and hESCs were used as controls for both experiments and analyses. Peak calling analysis for all samples using FDR< 0.01 yielded a total of 368,236 peaks for the ATAC-seq samples and 288,653 peaks for the ChIP-seq samples. First, we sought to understand whether the chromatin of GOKM and OSKM reprogrammable cells at D3 undergoes distinct remodeling towards the landscape of pluripotent cells or hTSCs. To address this question, we initially defined all ATAC-seq peaks that are specific to fibroblasts (75,225 peaks), hbdTSCs (49,101 peaks) and hESCs (70,086 peaks, Figure S3A). We then associated each peak to its closest neighboring gene and visualized the peaks and associated genes that define each cell type with scatterplots (Figure S3B-D, orange and light blue dots mark peaks that are associated with genes that are expressed in the corresponding cell type (i.e. hbdTSCs or in hESCs) according to RNA-seq data, while red and dark blue dots represent peaks in which no association with gene expression was identified). Low correlation coefficient (R^2^, 0.065-0.26) between the three samples, as well as peaks that associated with known cell type-specific genes (e.g. GATA3, TP63 and ELF5 in hbdTSCs, SOX2, SALL4 and NANOG in hESCs, POSTN and THY1 in fibroblasts) validated these sets of cell type-specific peaks (Figure S3B-D). Next, we used this information to identify newly remodeled ATAC-seq peaks from ‘GOKM D3’ and ‘OSKM D3’ (FDR< 0.05) samples by subtracting from these samples all the peaks that are shared exclusively with fibroblasts. This gave rise to 97,253 newly remodeled peaks that are specific for ‘GOKM D3’ and 52,121 newly remodeled peaks that are specific for ‘OSKM D3’, while 59,331 newly remodeled peaks were shared between the two combinations (Figure 3A). We overlapped ‘GOKM D3’ and ‘OSKM D3’ ATAC-seq peaks with hbdTSC-specific peaks or with hESC-specific peaks (Figure S3E-F). Interestingly, while ‘GOKM D3’ and ‘OSKM D3’ showed a comparable number of overlapping peaks with hESC-specific peaks (i.e. 3,168 peaks for ‘GOKM D3’ and 2,805 peaks for ‘OSKM D3’, Figure S3F), when hbdTSC-specific peaks were analyzed, GOKM D3 exhibited ~6-fold more overlapping peaks than OSKM D3 (i.e. 7,586 peaks for ‘GOKM D3’ and 1,306 peaks for ‘OSKM D3’, Figure S3E). These data suggest that already at day 3 of reprogramming, GOKM-induced cells show a bias toward remodeling of chromatin that is associated with hTSC state.

**Figure 3.**
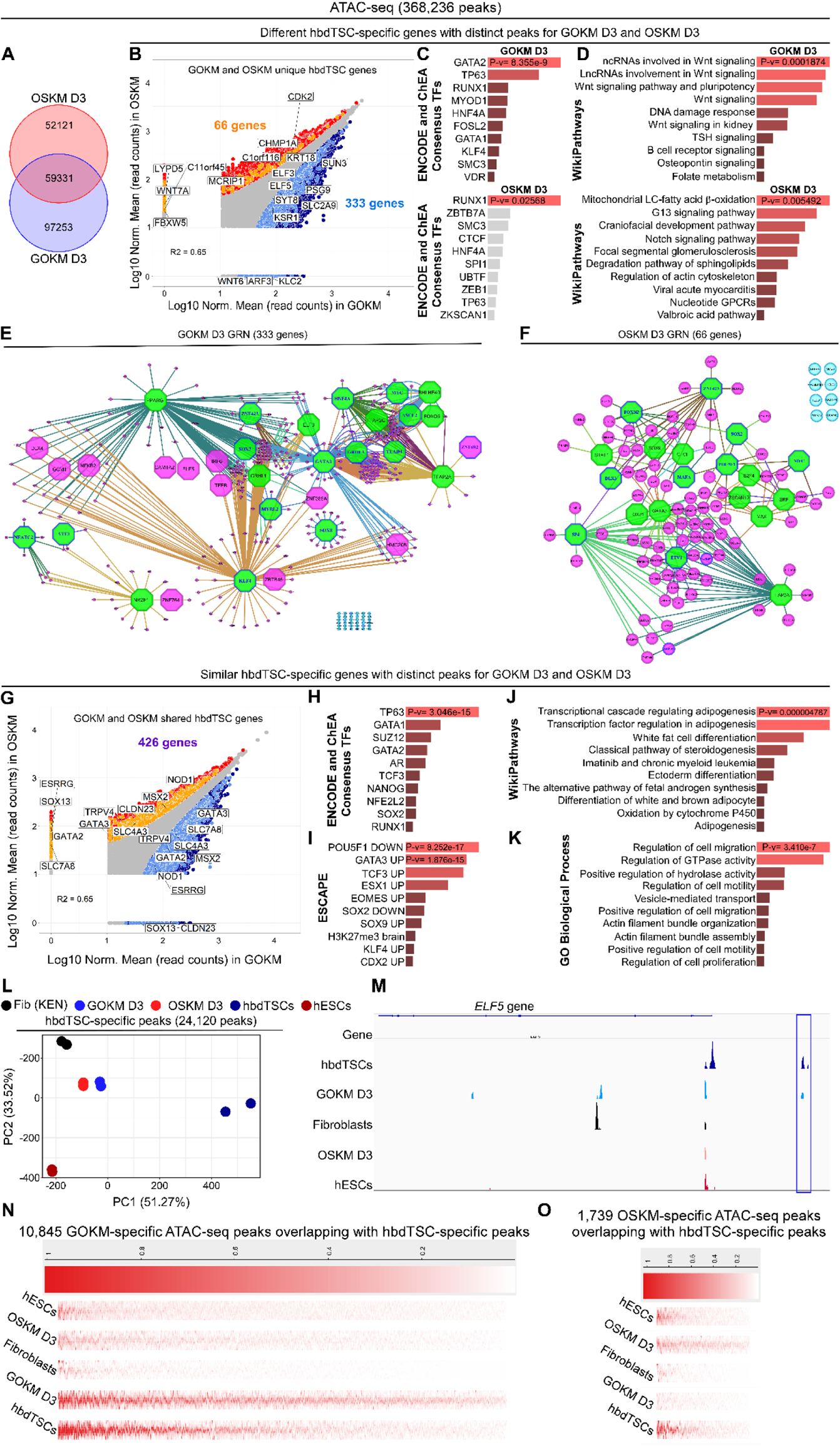
GOKM and OSKM each occupy the genome in a distinct manner and activate different placenta-related *loci* and pathways. To assess chromatin accessibility and activity of GOKM or OSKM early reprogrammable cells, GOKM or OSKM-transduced fibroblasts were exposed to dox for 72 hours and then collected and subjected to ATAC-seq analysis. **(A)** Venn diagram of GOKM D3 and OSKM D3 peaks after filtering of fibroblast-specific peaks FDR < 0.01. **(B)** A scatter plot of differentially accessible peaks between GOKM D3 and OSKM D3 with FDR< 0.05. Peaks that are specific to GOKM D3 or OSKM D3 are labeled with dark blue and red, respectively. Peaks associated with genes specific to day 3 of the GOKM reprogramming and are specific to the hbdTSC state are labeled with light blue, while peaks associated with genes specific to day 3 of the OSKM reprogramming and are specific to the hbdTSC state are labeled with orange. **(C-D)** Bar graphs showing the most enriched GO terms for 333 genes from (B) that were found to overlap between hbdTSC-specific genes and peaks that are associated with these genes in GOKM D3, and 66 genes from (B) that were found to overlap between hbdTSC-specific genes and peaks that are associated with these genes in OSKM D3 under the categories “ENCODE and ChEA Consensus TFs” (C) and “WikiPathways” (D), using EnrichR. **(E-F)** The TF genes regulatory network of 333 genes (E), and 66 genes (F) constructed by iRegulon plugin tool in Cytoscape where a transcription factor is considered if FDR< 0.05 and Network Enrichment Score (NES)> 2. Octagons represent genes that functions as a transcription factor. Green indicates transcription factors that functions as a pioneer regulator in the network, while pink is a regulated gene. Octagons that are pink represent transcription factors that are regulated by other transcription factors within the network. Genes that their expression levels were up-regulated significantly in day 3 of the reprogramming process are labeled with blue color. Genes with no significant association to transcription factors are marked by light blue. **(G)** Scatter plot of differentially accessible peaks between GOKM D3 and OSKM D3 with FDR< 0.05. Peaks that are specific to GOKM or OSKM are labeled by dark blue and red respectively. Peaks associated with genes that are shared between the hbdTSC state and in day 3 of the GOKM and OSKM reprogramming are labeled with light blue, or with light orange respectively. 426 genes are shared between the two reprogramming systems with different chromatin accessibility. **(H-K)** GO terms for 426 genes in (G) under the categories “ENCODE and ChEA Consensus TFs” (H), “WikiPathways” (J), Embryonic Stem Cell Atlas from Pluripotency Evidence “ESCAPE” (I) and GO biological process (K). **(L)** PCA analysis of 5 different groups across 24,120 accessibility regions that are specific to hbdTSCs. A clear separation between the two groups is observed, putting GOKM D3 in a closer distance toward the ‘hbdTSCs’ sample. **(M)** Normalized ATAC-seq profiles at the *ELF5 locus*. Profiles represent the union of all biological replicates for each group. **(N)** Heatmap of 10,845 open ATAC-seq GOKM-specific peaks overlapped solely with hbdTSCs peaks but not with OSKM (FDR< 0.05 and LFC> 1.5). **(O)** Heatmap of 1,739 open ATAC-seq OSKM-specific peaks overlapped solely with hbdTSCs peaks but not with GOKM (FDR< 0.05 and LFC> 1.5).

We then focused on the relation between the hbdTSC-specific ATAC-seq peak set and the newly remodeled peaks in ‘GOKM D3’ and ‘OSKM D3’. First, we associated GOKM D3 and OSKM D3 peaks to their neighboring genes and created scatterplots of the peaks which are unique to either GOKM D3 or OSKM D3. In these, we marked the genes that are expressed in hbdTSCs according to RNAseq data (Figures 3 B and G). This analysis revealed two groups of genes: (i) genes that are associated solely with GOKM peaks (333 genes) or genes that are associated solely with OSKM peaks (66 genes, Figure 3B), and (ii) genes that are associated with both ‘GOKM D3’ and ‘OSKM D3’ peaks (426 genes), with each factor combination generating peaks at a different genomic region along the gene’s *locus* (Figure 3G). For the first group, transcription factor binding site analysis revealed that GOKM remodel the chromatin of genes that are mainly regulated by GATA2 and TP63, transcription factors implicated in the TSC state ^1^, while OSKM remodel the chromatin of genes that are predominately regulated by the trophoblast differentiation factor RUNX1 (^31^, Figure 3C). GO term enrichment analysis identified WNT and DNA damage response as the major pathways for GOKM-regulated genes, while mitochondrial long-chain fatty acid β oxidation, G13 and NOTCH signaling were identified as the major pathways for OSKM-regulated genes (Figure 3D).

STRING and iRegulon analysis of the 333 GOKM-regulated genes (Figure 3E) and the 66 OSKM-regulated genes (Figure 3F) identified two distinct GRNs with 13 key hTSC transcription factors being expressed already at day 3 of GOKM transduction (blue framed octagons), and 7 hTSC transcription factors, along with OCT4 and SOX2 transgenes, in OSKM-transduced cells (Figure 3E-F). ‘GOKM D3’ GRN includes known hTSC state regulators such as TFAP2C, TEAD4, ASCL2, KLF4 and GRHL3, while ‘OSKM D3’ GRN is comprised of OCT4, SOX2, ETV1 and ZNF423. Both GRNs contain the two hTSC key transcription factors GATA3 and TFAP2A, as well as MYC (Figure 3E-F).

For the second group, transcription factor binding site and gene perturbation analyses revealed that both GOKM and OSKM remodel the chromatin of genes that are mainly regulated by TP63 and GATA1 and are affected when GATA3, TCF3 or EOMES are upregulated or when OCT4 and SOX2 are downregulated in hESCs (Figure 3H-I). GO term enrichment analysis identified adipogenesis and cell migration and motility as the major pathways that are regulated by these genes (Figure 3J-K).

Next, we focused on the relation between the hESC-specific ATAC-seq peak set and the newly remodeled peaks in ‘GOKM D3’ and ‘OSKM D3’. We again created scatterplots of the peaks which are unique to either ‘GOKM D3’ or ‘OSKM D3’, and this time we marked the genes that are expressed in hESCs according to RNAseq data (Figure S3G). This analysis identified 299 genes that are associated exclusively with GOKM peaks and 65 genes that are associated exclusively with OSKM peaks (Figure S3G). Transcription factor binding site analysis revealed that GOKM remodel the chromatin of genes that are mainly regulated by the transcription repressor REST while OSKM remodel the chromatin of genes that are mainly regulated by SUZ12 and KLF4 (Figure S3H). GO term enrichment analysis identified DNA replication and WNT signaling as the major pathways for GOKM-regulated genes, while focal adhesion-PI3K-Akt-mTOR signaling as the major pathway for OSKM-regulated genes (Figure S3I). We performed PCA using the most differential peaks across all samples (FDR < 1X10-7), while analyzing peaks that are either specific to (i) GOKM D3 and OSKM D3, (ii) hbdTSCs, or (iii) hESCs. In agreement with the previous results, GOKM demonstrated higher pioneer activity at day 3 of reprogramming compared to OSKM (Figure S3J). PCA performed on hbdTSC-specific peaks (Figure 3L) and hESC-specific peaks (Figure S3K) demonstrate that GOKM is superior to OSKM in remodeling the chromatin of the hTSC state (Figure 3L and two genomic loci representatives, *ELF5* and *TET3*, in Figures 3M and S3L), but not in remodeling the chromatin of the pluripotent state (Figure S3K).

Given that many pivotal stemness genes are shared between hESCs and hbdTSCs (e.g. LIN28A, FGF4, NR6A1, ZFP42, DPPA2, TET3, MYBL2), we next hypothesized that OSKM-specific peaks that are shared with hbdTSCs might be also shared with hESCs. To address this question, we took all peaks of hbdTSCs and subtracted only the peaks that overlapped with parental fibroblasts (FDR<0.05 and log fold change (LFC)>1.5). This gave rise to 126,125 peaks which we subsequently overlapped with either GOKM-specific peaks or with OSKM-specific peaks. In support of our hypothesis, the 10,845 GOKM-specific peaks overlapped solely with hbdTSC peaks but not with OSKM D3, parental fibroblasts or hESCs. In contrast, the 1,739 OSKM-specific peaks overlapped with both hbdTSCs and hESCs but not with GOKM D3 or parental fibroblasts (Figure 3N-O). These results imply that while the peaks shared between GOKM D3 and hbdTSCs reflect remodeling of the chromatin towards the hTSC state, those shared between hbdTSCs and OSKM D3 are mostly a result of overlap between chromatin which is remodeled also in pluripotency.

We then analyzed the ChIP-seq data for the histone mark H3K4me2. As was described for the ATAC-seq, we initially defined sets of peaks for each cell type and visualized the peaks and their associated genes using Venn diagrams and scatter plots (Figure S4A-D). We then extracted all the newly marked peaks from ‘GOKM D3’ and ‘OSKM D3’ samples by subtracting the peaks that are shared with fibroblasts. This resulted in 53,564 newly H3K4me2-marked peaks that are specific for ‘GOKM D3’ and 40,167 newly H3K4me2-marked peaks that are specific for ‘OSKM D3’, while 49,447 newly H3K4me2-marked peaks were shared between the two factor combinations (Figure S4E). We overlapped ‘GOKM D3’ and ‘OSKM D3’ peaks with hbdTSC-specific peaks or with hESC-specific peaks (Figure S4F-G). In agreement with the ATAC-seq analysis, GOKM D3 exhibited higher chromatin remodeling activity in hbdTSC *loci*, yielding ~4-fold more overlapping peaks with the hbdTSC-specific peak set when compared to OSKM D3 (i.e. 5,568 peaks for ‘GOKM D3’ and 1,532 peaks for ‘OSKM D3’, Figure S4F), while exhibiting a less pronounced difference (~2 fold more) with hESC-specific peaks (i.e. 4,627 peaks for ‘GOKM D3’ and 2,395 peaks for ‘OSKM D3’, Figure S4G).

We then examined the relation between the hbdTSC-specific H3K4me2 peak set and the newly marked peaks in GOKM D3 and OSKM D3. We created scatterplots of the peaks which are unique to either GOKM D3 or OSKM D3, and marked the genes that are expressed in hESCs according to RNAseq data (Figure S4H-I). This analysis too revealed two groups of genes: (i) genes that are associated solely with GOKM peaks (183 genes) and genes that are associated solely with OSKM peaks (161 genes, Figure S4H), and (ii) genes that are associated with both ‘GOKM D3’ and ‘OSKM D3’ peaks (225 genes) in which each factor combination generated peaks at a unique genomic region along the gene’s *locus* (Figure S4I).

Transcription factor binding site analysis for the first group revealed that while GOKM deposit the H3K4me2 histone mark on genes that are mainly regulated by GATA2 and GATA1, and to a lesser extent TP63, OSKM do so mainly via TP63, and to a lesser extent, by GATA2 and GATA1 (Figure S4J). GO term enrichment analysis identified Glucocorticoid metabolism as the major pathway of for GOKM-regulated genes, while WNT and NOTCH signaling as the major pathways for OSKM-regulated genes (Figure S4K). For the second group, transcription factor binding site and gene perturbation analyses revealed that both GOKM and OSKM remodel the chromatin of genes that are mainly regulated by TP63, AR and SUZ12 and are affected when the TSC key factors CDX2 or GATA3 are upregulated or when the pluripotency factors OCT4 and PRDM14 are downregulated in hESCs (Figure S4L-M). Comparable to the ATAC-seq analysis, GO term enrichment analysis identified adipogenesis, WNT signaling, GTPase activity and cell migration as the major pathways that are regulated by these genes (Figure S4N-O).

Finally, and in accordance with the ATAC-seq, a PCA plot generated using the most differential peaks (FDR < 1X10^−7^) between GOKM D3 and OSKM D3 and overlapped with hbdTSC-specific peaks again demonstrated that GOKM is superior in depositing the histone mark H3K4me2 on TSC-related *loci* compared to OSKM (Figure S4P).

Taken together, our results suggest that the GOKM and OSKM transcription factor combinations each remodel the chromatin in a unique manner at early stages of reprogramming, and that GOKM is more efficient in remodeling hTSC-specific *loci*. Focusing on areas shared with hbdTSCs, while GOKM alter the chromatin of genes responsible for hTSC stemness pathways such as WNT, DNA damage response and DNA replication, OSKM alter the chromatin of genes responsible for trophoblast differentiation pathways such as NOTCH, G13 and PI3K-mTOR-AKT signaling ^32–34^. Moreover, these data also shed light on the mechanism by which OSKM is capable of generating hiTSCs, albeit with a lower efficiency than GOKM.

### The reprogramming process toward hiTSCs does not induce genomic aberrations

Next, we asked whether the reprogramming process toward hiTSCs is prone to genomic aberrations. To that end, we subjected two hbdTSC lines, hbdTSC#2 and hbdTSC#9, and four hiTSC clones, hiTSC#4, hiTSC#11, hiTSC#2 and hiTSC#1, to sensitive karyotyping measurements using an Affymetrix CytoScan 750K array. Thorough analysis revealed that 50% of all colonies from both origins (i.e. hbdTSCs and hiTSCs) harbor an intact karyotype. The other 50% of the colonies exhibited few aberrations in a small fraction of the cells (Figure S5A). These results indicate that hiTSC colonies with an intact karyotype can be isolated and grown in culture and that the reprogramming process in itself does not facilitate genomic instability. However, the results do suggest that like in the mouse, hTSC cells have intrinsic tendency for genomic instability and that hTSC culture conditions should be optimized, as prolonged culture period might sensitize the cells for genomic aberrations, similarly to human ESCs/iPSCs ^35^. It is important to note that recent findings have highlighted the abundance of genomic aberrations and mosaicism in endogenous trophoblastic tissue as well ^36^.

### hiTSCs exhibit an extensive DNA de/methylation rewiring toward hTSC state

Given that the hiTSC gene expression profile is highly similar to that of hbdTSCs, we next wished to understand whether the epigenetic landscape of hiTSCs and hbdTSCs is correspondingly equivalent. DNA methylation is an epigenetic property that has been shown to be modified at late stages of OSKM reprogramming to iPSCs ^37^. To examine whether the DNA methylation landscape of hiTSCs is equivalent to that of hbdTSCs, we subjected five hiTSC clones (hiTSC#1, hiTSC#2, hiTSC#4, hiTSC#11 and hiTSC#16) to reduced representation bisulfite sequencing (RRBS) analysis, a method which increases depth of sequencing by focusing on genomic regions that are enriched for CpG content. Two hbdTSC lines, hbdTSC#2 and hbdTSC#9, two primary fibroblast lines (KEN and GM2) and hESCs were used as positive and negative controls, respectively.

Methylation analysis revealed 62,184 differentially methylated regions (DMRs) between fibroblasts and hbdTSC lines, 20,333 of which are hypomethylated in fibroblasts and hypermethylated in the hbdTSC lines, while the other 41,851 DMRs are hypermethylated in fibroblasts and hypomethylated in the hbdTSCs. Notably, analysis of the methylation landscape of the five hiTSC clones revealed robust de novo methylation in all five hiTSC clones, with a very small fraction of 1418 DMRs exhibiting partial de novo methylation, with their associated neighboring genes showing no significant association to hTSCs, hESCs or parental fibroblasts using GREAT and EnrichR (Figure 4A). In contrast, the 41,851 comparatively hypomethylated DMRs showed less efficient demethylation activity. Approximately one-third of the tiles, which display significant association to placental *loci*, demonstrated complete hypomethylation in all hiTSC colonies, while the remainder of the tiles exhibited only partial demethylation with high levels of variation between different hiTSC colonies (Figure 4B). Of note, one hiTSC colony, hiTSC#11, which was derived from the PCS201 fibroblast line, clustered closer to the two hbdTSC lines than to other hiTSC clones, which were derived from KEN (hiTSC#1, hiTSC#2, hiTSC#4) or GM2 (hiTSC#16) fibroblast lines (Figure 4B).

**Figure 4.**
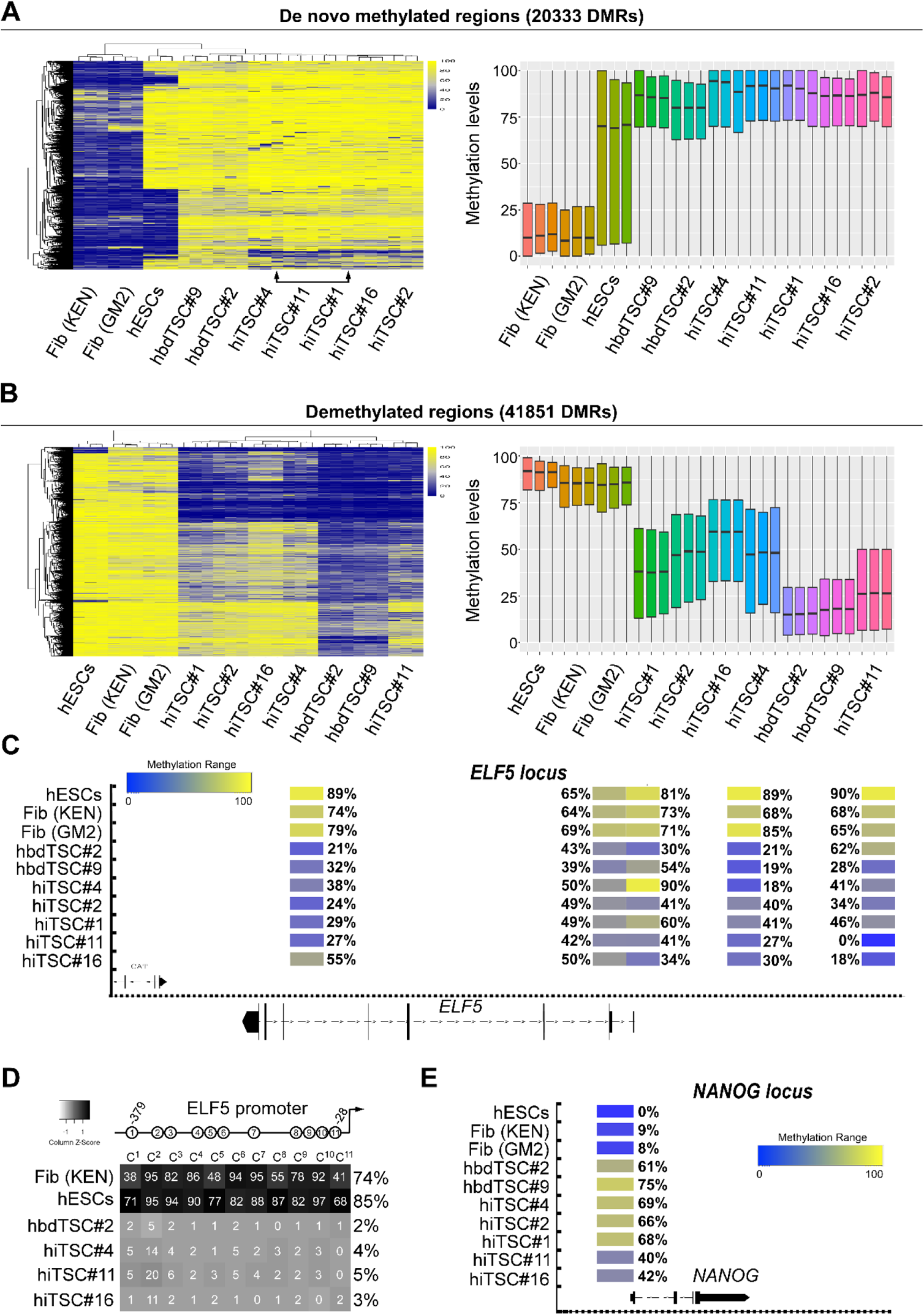
RRBS analysis demonstrates trophoblast-specific changes in methylation in hiTSCs. DNA Methylation analysis of three biological replicates of fibroblasts (KEN and GM2), hESCs, two hbdTSCs lines, hbdTSC#2 and hbdTSC#9, and five hiTSC clones, hiTSC#1, hiTSC#2, hiTSC#4, hiTSC#11 and hiTSC#16 as assessed by RRBS. Analysis of CpG methylation ratio with sequencing depth of at least 10 reads per CpG was computed, based on 100bp tiles. **(A, Left)** Heatmap showing 20333 differentially methylated regions (DMRs) that are hypomethylated in fibroblasts and hypermethylated in hbdTSCs with a methylation difference above 50%. hiTSCs are shown to have successfully acquired methylation patterns comparable to hbdTSCs. **(A, right)** Boxplot showing the average methylation levels for each biological sample. Bottom and top of the box are the 25th and 75th percentile (the lower and upper quartiles, respectively), and the band near the middle of the box is the 50th percentile (the median). Whiskers represent the highest and lowest values. **(B, left**) Heatmap showing 41851 DMRs that are hypermethylated in fibroblasts and hypomethylated in hbdTSCs with a methylation difference above 50%. Note that while some of the regions did not undergo a complete demethylation and that different hiTSC clones demonstrate different degree of demethylation, the overall demethylation landscape of all hiTSCs is more similar to hbdTSCs than fibroblasts or hESCs as indicated by the depicted dendrogram. **(B, right)** Box plot showing mean methylation pattern between fibroblasts and hbdTSCs colonies. Bottom and top of the box are the 25th and 75th percentile (the lower and upper quartiles, respectively), and the band near the middle of the box is the 50th percentile (the median). Whiskers represent the highest and lowest values. **(C)** Integrated genome browser capture of the methylation levels of five tiles that reside within the *ELF5 locus* in hESCs, fibroblast (KEN and GM2), hbdTSCs (hbdTSC#2 and hbdTSC#9), and five hiTSC clones, as assessed by RRBS. **(D**) Graph depicting average methylation levels of 11 CG within the proximal *ELF5* promoter, in the indicated samples, as assessed by targeted bisulfite sequencing using MiSeq 2×150 bp paired end run **(E)** Integrated genome browser capture of the methylation levels of one tile of the *NANOG locus,* in the indicated samples, as assessed by RRBS.

These results suggest that demethylation is less rigorous in hiTSC reprogramming, but may also imply that the background, sex and the age of the parental fibroblasts may play a role in the efficiency of the demethylation process (i.e. GM2-derived hiTSC#16 clone that showed the lowest DNA demethylation capabilities was derived from fibroblasts isolated from adult female, compared to KEN and PSC, which are both HFF lines). Importantly, although the demethylation process is not optimal in hiTSCs, the overall methylation landscape of hiTSC clones clustered closely to hbdTSCs and far from hESC and fibroblast controls in both de novo methylated and demethylated DMRs (Figure 4A-B).

Several gatekeeper genes have been found to remain methylated during mouse ESC transdifferentiation toward TSC fate, producing cells which are TS-like but do not acquire complete TSC identity ^12^. One of these gatekeeper genes is *ELF5*, the demethylation thereof is considered an important criterion for human trophoblast cell identity ^23^. Thus, we examined whether the *ELF5* locus underwent demethylation in hiTSCs. Analysis of the RRBS data showed five DMRs in the *ELF5 locus*, demonstrating an overall equivalent pattern of hypomethylation between the two hbdTSC clones and all hiTSC clones (Figure 4C). In agreement with the RRBS results, targeted direct amplification and next generation sequencing of the proximal *ELF5* promoter region after bisulfite conversion demonstrated vigorous demethylation in all 11 CpG promoter sites in both hbdTSCs and hiTSCs, but not in hESC and fibroblast controls (Figure 4D).

We next examined the methylation levels of the pluripotency-specific locus *NANOG*, which is hypermethylated in mouse TSCs ^10, 13^. Similar to the mouse, the only DMR from the RRBS data that received coverage in this locus was completely hypomethylated in hESCs and to a lesser extent in fibroblasts, but equivalently methylated in both hbdTSCs and hiTSCs (Figure 4E).

Taken together, these data suggest that DNA methylation is largely rewired to the hTSC state in the stable hiTSCs, but also suggest that improved reprogramming conditions need to be developed to induce a more robust demethylation.

### hiTSCs differentiate into all major trophoblast cell types with variation in EVT differentiation

hTSCs have the ability to differentiate into multinucleated syncytiotrophoblast (STs) and extravillous trophoblasts (EVTs) ^1^. Thus, our next goal was to examine whether hiTSCs have the potential to differentiate into these various trophoblast subtypes.

We performed directed differentiation into STs and EVTs using previously published protocols ^1^. Initially, we differentiated hbdTSCs and three hiTSC clones (hiTSC#4, hiTSC#11 and hiTSC#16) into STs and collected samples at days 2 and 6 of differentiation. qPCR analysis for ST markers such as *CSH1*, *GCM1, SDC1, CGB, PSG1, CHSY1 and ERVFRD-1* ^1, 23^ showed robust induction of ST markers in hiTSCs, equivalent to in hbdTSCs (Figures 5A and S5B). Of note, ERVFRD-1, which is an endogenous retroviral gene expressed by cytotrophoblastic ST-precursor cells and orchestrates the fusion event early in the syncytialization process ^38^, showed a rapid and transient upregulation on day 2 of differentiation (Figure 5A). Bright field images and immunostaining for the pan trophoblast KRT7, the epithelial marker CDH1 and DAPI showed clear formation of large KRT7-positive multinucleated cells after 6 days of differentiation in both hbdTSCs and hiTSCs (Figure 5B). As expected, while the undifferentiated hiTSCs stained positive for CDH1 (Figure 1E), staining in multinucleated STs was significantly reduced. CSH1 and SDC1-positive three-dimensional ST structures were observed in all hiTSC clones and the hbdTSC#2 positive control (Figure 5B). These data indicate that hiTSCs are capable of differentiating into STs similarly to hbdTSCs.

**Figure 5.**
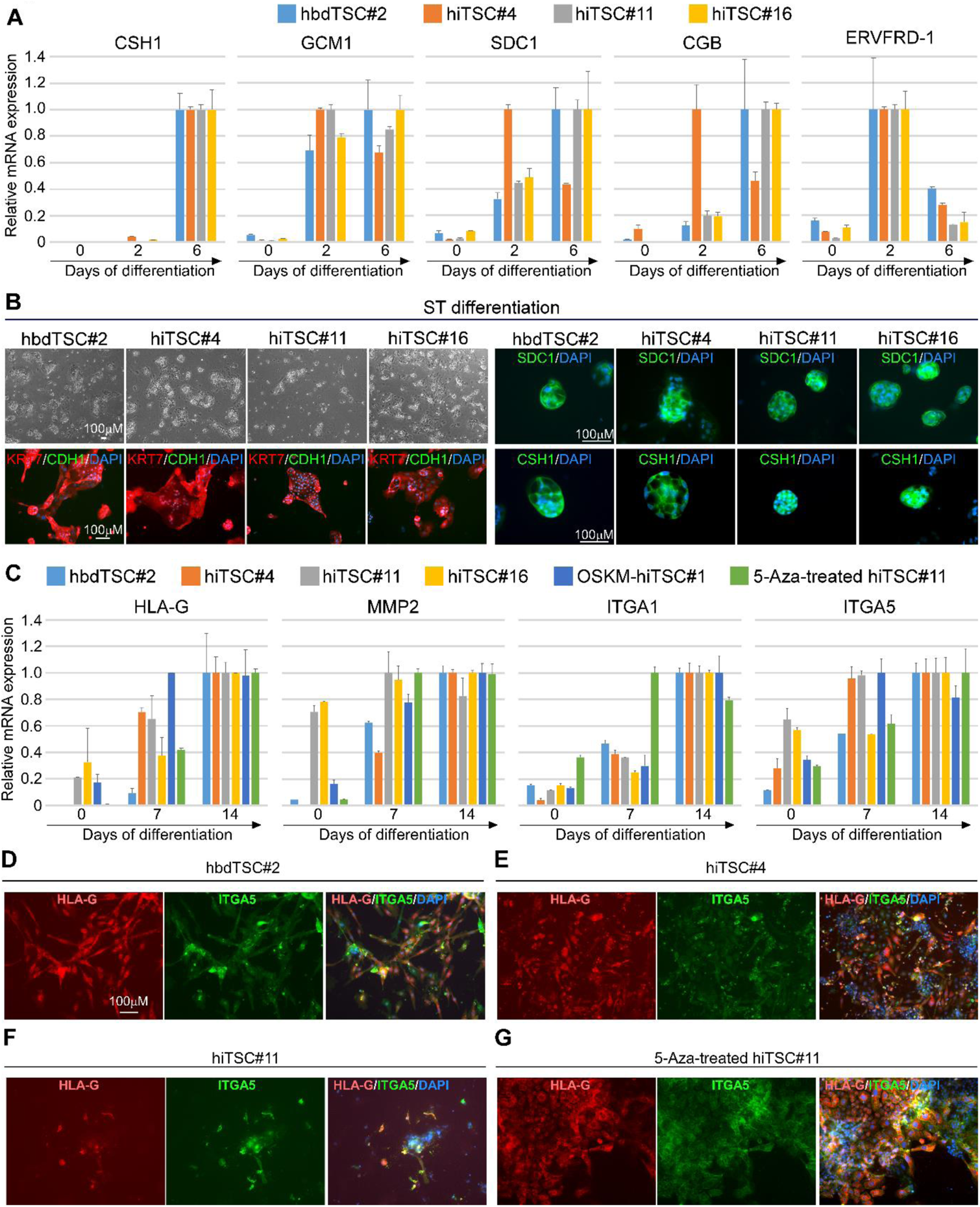
hiTSCs differentiate into multinucleated ST and EVT cells. hbdTSC#2, hiTSC#4, hiTSC#11 and hiTSC#16 were induced to differentiate into STs using previous protocol ^1^. **(A)** qPCR analysis of relative mRNA levels of ST-specific markers *CSH1*, *GCM1*, *SDC1*, *CGB* and *ERVFRD-1*, for the indicated samples, at days 0, 2 and 6 in medium for directed differentiation into ST (STM). The highest sample for each gene per colony was set to 1. Results were normalized to the mRNA levels of the housekeeping control gene *GAPDH* and are shown as fold change. Error bars indicate standard deviation between two duplicates. **(B)**. Bright field images (left top) and fluorescent images for the pan-trophoblast marker KRT7 and the epithelial marker CDH1 (left bottom), or for ST-specific markers SDC1 and CSH1 (right panel) following 6 days of directed differentiation to STs. DAPI staining was included to mark nuclei. Two and three dimensional multinucleated-positive cells are shown for each indicated clones. **(C)** qPCR analysis of relative mRNA levels of EVT-specific markers *HLA-G*, *MMP2*, *ITGA5* and *ITGA1* at days 0, 6 and 14 of directed differentiation into EVT. The highest sample for each gene per clone was set to 1. Results were normalized to the mRNA levels of the housekeeping control gene *GAPDH* and are shown as fold change. Bars indicate standard deviation between two duplicates. **(D-G)** Immunofluorescent staining for the EVT-specific markers HLA-G and ITGA5 and DAPI in PFA-fixated hbdTSC#2 (D), hiTSC#4 (E), hiTSC#11 (F) and 5-Aza-treated hiTSC#11 (F) following 14 days of EVT differentiation.

We next performed directed differentiation of hbdTSCs and hiTSCs into EVTs ^1^. Following seeding and cell attachment to the plate, cell aggregates formed in all tested clones. However, after 14 days of differentiation, the efficiency in the production of HLA-G-positive EVTs varied widely between clones (Figure 5C). While hbdTSC#2 and hiTSC#4 showed massive differentiation into EVTs as assessed by morphology, EVT gene expression (i.e *HLA-G*, *MMP2*, *ITGA5* and *ITGA1,* ^1, 23^) and staining for the EVT markers HLA-G and ITGA5 (Figure 5C-E), hiTSC#11 and hiTSC#16 as well as OSKM-hiTSC#1 demonstrated only a partial capability in producing HLA-G-positive cells (Figures 5C-F and S5C). In general, we observed two main morphologies of HLA-G-positive cells following 14 days of EVT differentiation; (i) spindle-shaped and (ii) small and migratory cells. Interestingly, while the spindle-shaped cells exhibited strong HLA-G staining and weak ITGA5 expression, the small and migratory cells stained strongly for both HLA-G and ITGA5 (Figure 5D-E). The appearance of multiple morphologies of EVTs in our experiments is in accordance with previous studies showing significant diversity between various EVT subtypes ^39^.

We sought to examine whether the partial DNA demethylation observed in some hiTSC clones or reactivation of viral vectors during differentiation might contribute to the inconsistency seen during EVT differentiation. To that end, we treated hiTSC#11 and hiTSC#16 with 5-aza-2′-deoxycytidine (5-Aza), a known demethylation agent, for two days and performed EVT differentiation. In parallel, we utilized a previously described non-integrating episomal reprogramming technique ^40^, replacing the hSOX2 with hGATA3, and reprogrammed the cells into hiTSCs.

Interestingly, one of two 5-Aza treated GOKM-hiTSC clones (i.e. hiTSC#11) restored its capability to differentiate into EVTs (Figure 5G) and 1 out of 3 examined episomal-derived hiTSC clones showed robust differentiation into EVTs (Figure S5D).

These results suggest that suboptimal demethylation can hinder the differentiation potential of the cells and that transgene integration most probably do not contribute to the inconsistency seen in EVT differentiation between hiTSC colonies. It is important to note that hiPSC clones also harbor various propensities for differentiation ^41^ and that this variability is an intrinsic property within any reprogrammed cells.

Taken together, these results suggest that some hiTSC clones hold a full differentiation potential, but also imply that the current methodology of hiTSC formation and differentiation protocols need to be optimized to unleash the full differentiation potential of the cells.

### hiTSCs form trophoblastic lesions following subcutaneous injection into NOD/SCID mice

When mouse TSCs/iTSCs are injected subcutaneously into nude mice, the cells differentiate into the various trophoblast subtypes and orchestrate robust invasion and endothelial cell recruitment to form transient hemorrhagic lesions. Conversely, when hbdTSCs are injected subcutaneously into non-obese diabetic (NOD)-severe combined immunodeficiency (SCID) mice, the cells form KRT7-positive trophoblastic lesions with little differentiation and blood vessel formation ^1^. To test whether hiTSCs are capable of forming similar trophoblastic lesions, we subcutaneously injected ~4×10^6^ cells from three hiTSC clones (hiTSC#4, hiTSC#11 and hiTSC#16) and one hbdTSC control line hbdTSC#2, into NOD/SCID male mice. Nine days later, when the lesions reached approximately 5mm in size (Figure 6A), the lesions and blood serum were extracted and examined. Using enzyme-linked immunosorbent assay (ELISA), we quantified the level of the human pregnancy hormone hCG in the serum of the injected male mice. When fibroblasts were injected, the serum levels of hCG were undetectable, while serum levels following injection of hiTSC and hbdTSC clones was in the range of 40-130 mlU/ml (Figure 6B). Immunohistochemical staining showed that all lesions were KRT7-positive and that small regions of cells had differentiated into EVTs and STs (Figure S6A), similarly to previously published findings ^1^. These results imply that hiTSCs can generate trophoblastic lesions in NOD/SCID mice that are similar in their characteristics to lesions that are formed by hbdTSCs.

**Figure 6.**
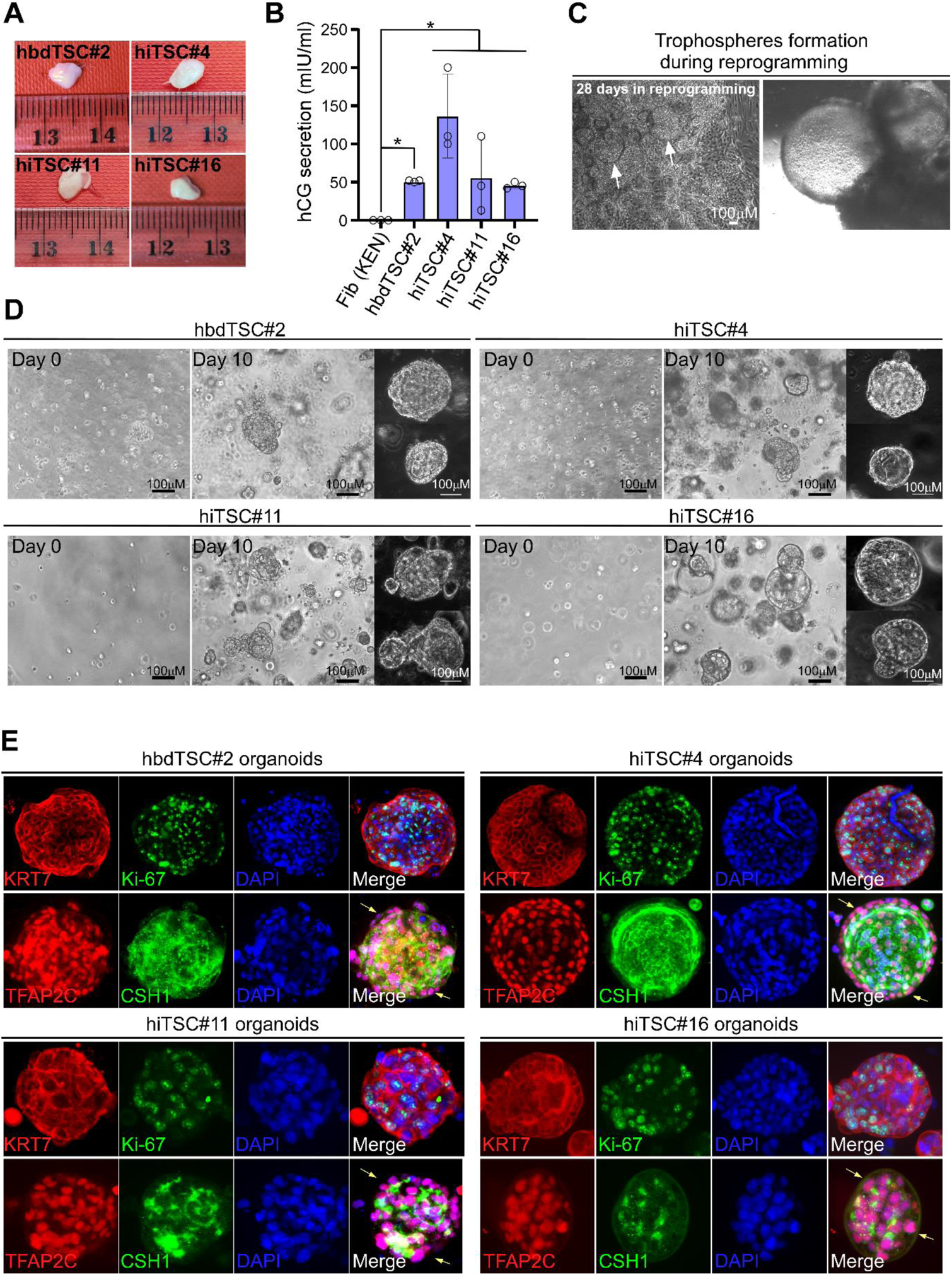
hiTSCs form trophoblastic lesions in NOD/SCID mice and functional organoids in matrigel. **(A)** Image of the lesions that were extracted from NOD/SCID mice following ~9 days of subcutaneous injection of ~4×10^6^ cells from hbdTSC#2, hiTSC#4, hiTSC#11 and hiTSC#16 lines. **(B)** Graph showing the concentration of hCG secretion in the serum of injected NOD/SCID male mice with the indicated cells. Approximately 500μl of serum was collected from each mouse and used for hCG detection, using hCG ELISA kit (Alpco). Error bars indicate standard deviation between 3 different mice. *p-value< 0.05 as calculated by GraphPad Prism using Student’s t-test. **(C)** Bright field images of trophospheres in a reprogramming plate at day 28 of dox treatment following GOKM infection. **(D)** Bright field images of hbdTSC#2, hiTSC#4, hiTSC#11 and hiTSC#16 at day 0 and 10 of organoid formation protocol. Note that by day 10 of the protocol, large and 3-dimantional organoid structures were generated in each clone. **(E)** Spinning disk confocal images of organoids from (D), following immunofluorescence staining for pan-trophoblast marker KRT7, proliferative cell marker Ki-67, TFAP2C and CSH1, with DAPI nuclear staining. Yellow arrows indicate areas of undifferentiated cells.

### hiTSCs form functional trophoblast organoids

Recently, two trophoblast organoid systems have been developed and described ^42, 43^. These studies demonstrated the capability of first trimester villous CTB cells to form three-dimensional structures which contain both proliferating stem cells and differentiated cells. In-depth examination of the two systems revealed that many characteristics of the early developmental program of the human placenta are present in these organoid platforms ^42, 43^.

Thus, we next asked whether hiTSCs harbor similar potential to form trophoblastic spheres and functional organoids. Initially we observed, albeit occasionally, regions within the reprogramming plates which generated trophoblastic spheres (Figure 6C), suggesting that these cells are capable of forming three-dimensional structures. Notably, these trophospheres were recently suggested to mark naïve hTSC (i.e. similar to pre-implantation TE, ^7^). To examine whether isolated hTSC clones have a similar capacity, we employed a previously published protocol for trophoblast organoid formation ^42^ on single cells and tested organoid formation. hiTSC#4, hiTSC#11, hiTSC#16 and control hbdTSC#2 were trypsinized and seeded as single cells inside a droplet of matrigel and allowed to grow for 10 days. As shown for villous CTBs ^42, 43^, both hbdTSCs and hiTSC clones were capable of forming three-dimensional structures within few days of culture (Figure 6D). Immunostaining for the pan-trophoblast marker KRT7 and the proliferation marker Ki-67 followed by confocal microscopy examination revealed KRT7-positive organoids with proliferating cells, suggesting that undifferentiated cells were still present within the organoids after 10 days of culture (Figure 6E). Immunostaining for TFAP2C and CSH1 validated an outer layer of cytotrophoblasts with a core of STs, displaying the inverted placental villous structure as previously described ^42, 43^. Taken together, these data demonstrate that hiTSCs can form functional organoids that are similar to their hbdTSC and villous CTB counterparts.

### The reprogramming process into hiTSCs bypasses a pluripotent stage

Given that OSKM can generate both hiPSCs and hiTSCs ^3, 9, 44^ and since we show here that GATA3 can replace SOX2 in generating hiTSCs, we next asked whether OCT4, KLF4 and MYC (OKM) are sufficient to generate hiTSCs. Figures S1C and S7A-E demonstrate that the OKM combination is, in principle, incapable of producing neither hiTSCs nor iPSCs as assessed by colony number and expression of hTSC and hiPSC gene markers in reprogramming plates (Figure S1C and S7A-E). Of note, very rarely and following multiple attempts, we surprisingly managed to isolate two hiPSC colonies, but not hiTSC colonies, with the OKM factors only (Figure S6B).

Additionally, since multiple components of the hTSC medium have been shown to enhance mouse and human reprogramming to iPSC ^45, 46^, we tested the capability of GOKM to generate hiPSCs. The lack of hiPSC marker expression in hiTSC reprogramming plates hints that the hTSC medium does not support pluripotent cells (Figure S7D). Nevertheless, we transduced fibroblasts with the GOKM factors and conducted reprogramming with an established hiPSC reprogramming protocol. No hiPSC colonies emerged. We next repeated this experiment, but instead of using the hiPSC reprogramming protocol, we began with the TSC reprogramming protocol and upon dox withdrawal we changed the medium to hESC-supportive medium. Following three independent reprogramming experiments, we observed the formation of only two hiPSC colonies that were positive to hESC markers and integrated all four GOKM transgenes (Figures S6B and S7F). According to the transgene integration analysis, it is evident that the GOKM-iPSC colonies integrated higher levels of MYC and lower levels of KLF4 in comparison with the GOKM-iTSC colonies (Figure S7F). The importance of factor stoichiometry in determining cell identity was recently shown by our group when one combination of five transcription factors was able to induce three different cell types of the mouse preimplantation embryo depending on various transgene levels ^11^.

In order to further scrutinize the formation of hiPSCs during GOKM reprogramming, we sorted for pluripotency cell-surface marker TRA-1-60 positive cells at various time points during GOKM reprogramming, then seeded them and counted the hiPSC colonies which emerged (Figure S7G). We chose to sort even very weakly positive cells as so not to omit any potential hiPSCs at the cost of a higher false-positive sorting rate. Although hiPSC colonies emerged after seeding sorted cells from OSKM reprogramming and hiPSC positive controls, no hiPSC colonies emerged in plates seeded with cells sorted from GOKM reprogramming in various time points in the iTSC reprogramming protocol, following GOKM reprogramming with hiPSC protocol or after switching to hESC-supportive medium after dox withdrawal following hiTSC protocol (Figure S7G).

Although the emergence of hiPSC colonies with GOKM is evidently extremely rare, it prompts further investigation of whether GOKM-derived hiTSCs undergo a stage of transient pluripotency. To address whether pluripotency is a requirement for hiTSC formation, we utilized a previously published lentiviral-vector constitutively expressing the CRISPR/Cas9 protein and gRNA, which was shown to produce a very high occurrence of indels ^47^. Using bulk infection, we generated a heterogeneous population of SOX2 knockout (KO) fibroblasts that also express the Tet-On system transactivator, M2rtTA (Figure 7A). Since pluripotency cannot be maintained without SOX2 ^48^, obtaining hiTSC colonies that are SOX2 KO with GOKM indicates that pluripotency is not required for achieving the hTSC state during reprogramming. SOX2 KO fibroblasts were reprogrammed into hiTSCs with GOKM and several hiTSC colonies were isolated and propagated. Careful examination of the resulting colonies revealed that 7 out of 7 examined colonies contained SOX2 indels (Figure 7B-D) and 4 out of 7 contained a bi-allelic deletion within the SOX2 coding region (Figure 7E). SOX2-KO hiTSC colonies exhibited normal morphology (Figure 7F) and comparable gene expression to WT hiTSCs and hbdTSCs (Figure S7H). To confirm functional KO of SOX2 in the cells, we reprogrammed SOX2 KO fibroblasts into hiPSCs with human OKM and mouse Sox2 vector. This approach allowed us to distinguish between the endogenous human SOX2 from the exogenous mouse Sox2 gene in the resulting colonies. We were able to generate hiPSC colonies with OKM and mouse Sox2, although following dox withdrawal many hiPSCs collapsed and the ones that retained pluripotency were KO-escapees, which harbored at least one functional allele of SOX2 as assessed by qPCR analysis (S7I-L). This was in stark contrast to hiTSCs that demonstrated full homozygous deletion in 4 out of 6 examined colonies (Figure S7I-L). To validate our results, we generated an additional KO fibroblast line in which both NANOG and PRDM14 were targeted with specific gRNAs (Figure 7G). In agreement with the SOX2 KO results, not only were GOKM capable of producing hiTSCs from these DKO fibroblasts, the efficiency of hiTSC formation was significantly improved, yielding 2.5 fold more hiTSCs colonies compared to wild type (WT) fibroblasts (Figure 7H). Furthermore, DKO fibroblasts showed significantly reduced capacity to reprogram into hiPSCs (Figure S7M). Sequencing of the gRNAs regions of seven DKO hiTSC colonies followed by chromatogram analysis revealed that 6/7 colonies contain homozygous indels, that change the frameshift of the gene, for both PRDM14 and NANOG, while 1/7 colonies contains homozygous indels for PRDM14 and a heterozygous indel for NANOG (Figure 7I).

**Figure 7.**
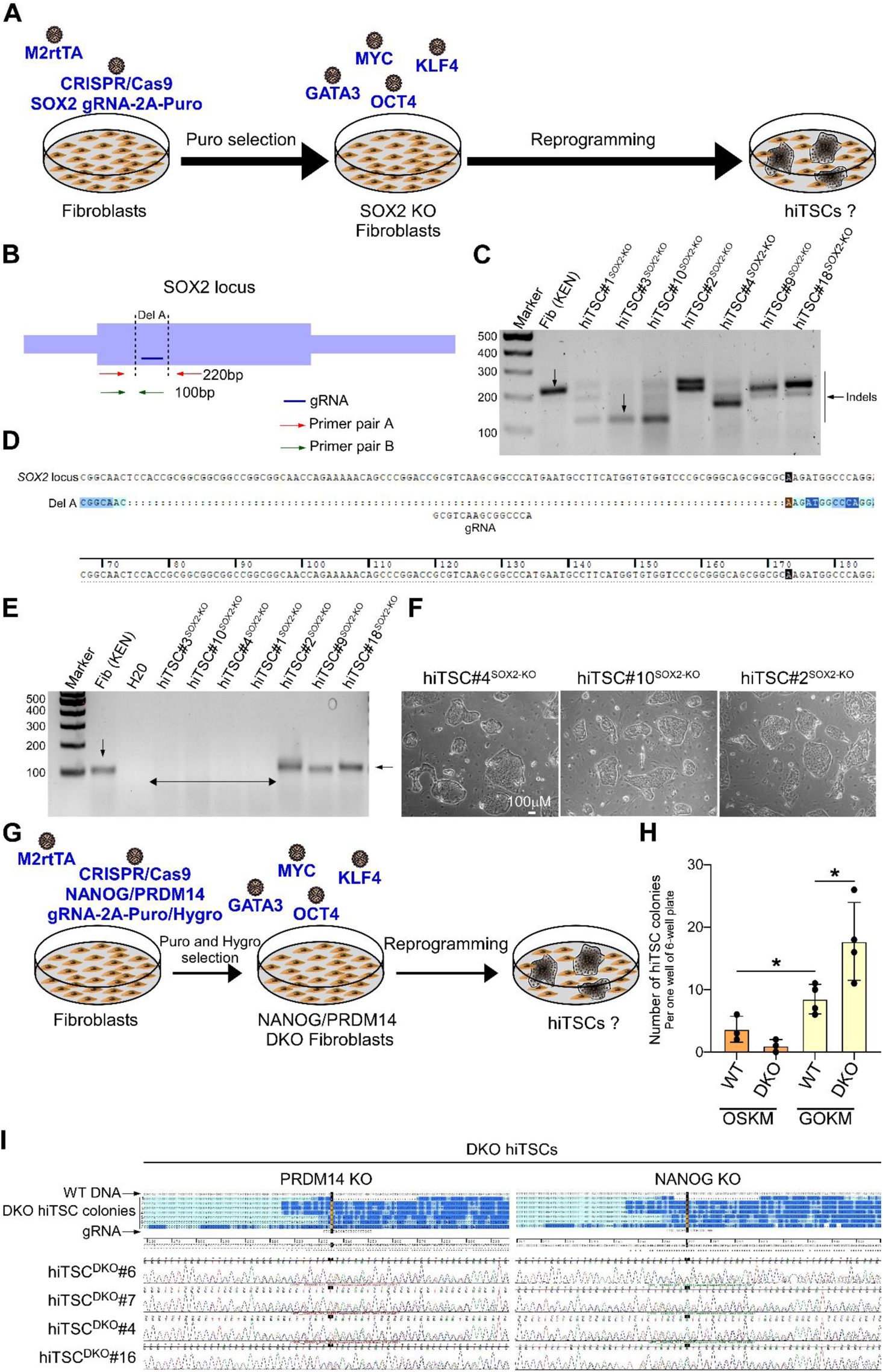
GOKM produce hiTSCs independently to SOX2 or PRDM14 and NANOG expression. **(A)** Schematic representation of knockout (KO) strategy of SOX2 in fibroblasts (KEN) and reprogramming into hiTSCs. Fibroblasts were first infected with a lentiviral vector inducing constitutive expression of Cas9 together with a guide RNA to the SOX2 gene coding region, as well as with M2rtTA, and selected with puro. Then, fibroblasts were reprogrammed as described to hiTSCs with GOKM. **(B)** Schematic representation of the *SOX2* gene, displaying the location of the gRNA used, as well as the primer pairs designed to examine the presence of indels. **(C)** DNA gel showing a WT band of 219bp within the *SOX2* coding region in WT HFFs and the same PCR reaction in seven independent hiTSC clones from fibroblasts which contained SOX2 indels. **(D)** A sequence alignment image of one indel event (i.e. Del A) within the *SOX2* locus in hiTSC#3^SOX2-KO^ using Sequencher software. **(E)** DNA gel showing a WT band of 100bp within the *SOX2* coding region in WT HFFs and the same PCR reaction in seven independent hiTSC clones from fibroblasts which contained SOX2 indels. Note, 4 out of 7 colonies show a bi-allelic deletion within SOX2 coding region (SOX2del/del). **(F)** Bright field images of three SOX2 KO hiTSC colonies. **(G)** Schematic representation of double knockout (DKO) strategy of *NANOG* and *PRDM14* in fibroblasts and reprogramming into hiTSCs. M2rtTA-transduced fibroblasts were co-infected with a lentiviral vector encoding for puromycin resistance and constitutive active Cas9 together with a gRNA against the *NANOG* coding region, as well as a lentiviral vector encoding for hygromycin resistance and constitutive active Cas9 together with a gRNA against the *PRDM14* coding region. Puromycin and hygromycin selected fibroblasts and WT controls were then reprogrammed as described to hiTSCs with GOKM. **(H)** Graph depicting the number of hiTSC colonies that emerged either in DKO fibroblasts or in WT controls following reprogramming with GOKM or OSKM. Note that while the loss of PRDM14 and NANOG facilitates the emergence of hiTSCs colonies with GOKM, it had no effect on the number of hiTSCs with OSKM. Error bars indicate standard deviation between 3-4 replicates. *p-value< 0.05 as calculated by GraphPad Prism using Student’s t-test. **(I)** Sequences alignment image of various indel events within the *NANOG* and *PRDM14 loci* in seven hiTSC^DKO^ clones using Sequencher software. gRNA sequences and Sanger chromatograms are shown for 4 hiTSC^DKO^ clones. Note that a significant enrichment for double KO events is evident in hiTSC clones that were derived from DKO fibroblasts.

Taken together, these data not only indicate that SOX2, NANOG and PRDM14 expression is dispensable for the formation of hiTSCs and that hiTSC reprogramming does not pass through a pluripotent state, but further hint that pluripotency-specific genes interfere with the acquisition of the hTSC state.

## DISCUSSION

Placental disorders such as preeclampsia and intra-uterine growth restriction are commonly detected at late stages of pregnancy, when proliferative villous CTBs are no longer available for isolation and exploration. Thus, developing a method to reprogram differentiated cells derived from disease-affected placenta or cord blood into functional TSCs is of vital importance for modeling and identifying potential risk factors for placental disorders, as well as for possible future cell-based therapy for supporting implantation in cases of recurrent miscarriages.

The generation of hiTSCs from naïve hiPSCs or following OSKM pluripotency reprogramming has been recently described ^3–9^, proposing one possible strategy of producing hiTSCs from mesenchymal cells. However, the production of hiTSCs from mesenchymal cells, independently of pluripotency or pluripotency factors, has not been shown before.

The direct conversion of fibroblasts into hiTSCs provides multiple advantages, both for basic research and for disease modelling approaches when compared to pluripotency-dependent hiTSC production. These include the following: (1) Direct conversion is capable of illuminating hTSC master regulators that are necessary for the induction of the hTSC state, while pluripotency-dependent protocols (i.e. OSKM reprogramming or human naïve ESC/iPSC transdifferentiation) mask those key master regulators. (2) Reprogramming to pluripotency in primed hESC conditions maintains normal methylation in *imprinted loci* while naïve pluripotency conditions have been shown to result in defective methylation in *imprinted loci* ^49^. As imprinting plays a pivotal role during the specification of the TE state, aberrant methylation at *imprinted loci* holds a significant limitation using these conditions. (3) In the same vein, naïve human pluripotency conditions induce global and non-specific DNA hypomethylation that may permanently erase important disease-related epigenetic defects that will mask the true driver/s of the disease. These epigenetic changes may be the result of intrinsic genetic tendencies or the effects of a specific intra-uterine environment. (4) In the mouse, the pluripotent state acts as a barrier for the production of fully functional hiTSCs by retaining methylation in TSC gatekeepers ^12^, while direct conversion produces miTSCs with normal methylation at gatekeeper *loci* ^10, 13^. (5) Residual pluripotent stem cells or those which are not fully differentiated have the potential to generate teratocarcinomas, raising concern when considering using such protocols for cell-based therapy.

Here, we demonstrate that transient expression of GATA3, OCT4, KLF4 and MYC (GOKM) is capable of directly converting male and female human fibroblasts into functional hiTSCs without acquiring a pluripotent state. We reveal that, although essential for mouse TSC circuitry ^50^, SOX2 is dispensable for the induction of the hTSC state and that GATA3 together with OCT4 are main drivers of hTSC identity. Though previous works have uncovered a central role for OCT4 in the establishment of the human trophoblast lineage ^20^, our work confirms that this role is independent of its known function as a master regulator of pluripotency.

We show that GOKM target the chromatin differently than OSKM and that GOKM reprogramming is specifically directed toward the hTSC state. In contrast, OSKM predominantly target regions that are shared between hESCs and hbdTSCs, suggesting an explanation as for how OSKM are also capable of generating hiTSCs. One interesting observation is that GOKM demonstrate a greater pioneer activity at early stages of reprogramming than OSKM as assessed by the higher number of regions that are remodeled and defined along the somatic genome. In accordance with these data, GOKM are capable of generating hiTSCs more efficiently than OSKM.

Methylation analysis revealed that the overwhelming majority of hbdTSC-specific methylated regions, compared to fibroblasts, underwent appropriate de novo methylation in the hiTSC clones. In contrast, a higher variation was seen between the various hiTSC clones in the hbdTSC-demethylated regions. Notwithstanding, one of the hiTSC clones clustered closer to hbdTSCs than to the other hiTSC clones, suggesting that near complete DNA methylation reprogramming is possible with GOKM reprogramming. Importantly, gate keepers such as *ELF5* that are abnormally methylated during mouse ESC-TSC transdifferentiation are demethylated properly in the reprogrammed hiTSCs, similarly to the mouse ^12, 23^.

It is important to note that appropriate rewiring of the methylation landscape during reprogramming is essential for both driving the reprogramming process and for acquiring the full epigenetic state of the targeted cells. Thus, we believe that improving demethylation capability by optimizing culture conditions and factor transduction will not only unleash the full functional potential of the cells, but will likely also increase the reprogramming efficiency. Devising simple methods such as gene expression markers to screen higher quality from lower quality reprogrammed colonies would also be helpful in obtaining optimal hiTSC lines.

Given recent literature showing the capacity to derive hiTSCs from naïve pluripotency conditions and reprogramming, and following the issues mentioned above regarding pluripotency and the hTSC state, it was important to confirm that GOKM hiTSCs do not rely on obtaining a transient pluripotent state. By knocking out SOX2 or NANOG/PRDM14 in fibroblasts, we show that not only is pluripotency not required for the formation of hiTSCs, but rather it likely hinders hiTSC production, as a greater number of colonies emerged in the KO fibroblast lines compared to WT fibroblasts.

Functional experiments validated hiTSC identity and demonstrated full developmental potential as assessed by capability to differentiate into STs and EVTs, form trophoblastic lesions in NOD/SCID mice and develop organoids in matrigel. We believe that the variations in EVT differentiation noted between different hiTSC colonies may be a result of the variation observed in the DNA methylation landscape of the cells. In support of this assumption, 5-Aza treatment facilitated EVT differentiation in some EVT-refractory treated clones.

Overall, we describe a system to produce fully functional hiTSCs from mesenchymal cells originating from either male or female, neonate or adult individuals. Though some calibration of culture and reprogramming conditions is likely needed, the advantages of the direct conversion approach are significant. We offer this reprogramming strategy as a valuable source to study diseases that are associated with pathological placental development.

## ACKNOWLEDGMENTS

Y.B. is supported by a gift from the Morningstar Foundation and Edward & Millie Carew-Shaw Distinguished Medical Faculty Award and research grants from the European Research Council (ERC, #676843), the Israeli Center of Research Excellence (I-CORE) program (Center #41/11), the Israel Science Foundation (ISF, #823/14), EMBO Young Investigator Programme (YIP), Kamin (#53776), Abisch-Frenkel Foundation (#15/H5), Alon foundation Scholar–Program for distinguished junior faculty, the American Society for Reproductive Medicine, DKFZ-MOST (177), MOST (88507) and Howard Hughes Medical Institute International Research Scholar (HHMI, #55008727). A.R. is supported by the UK-Israel BIRAX 030-5187 fellowship.

## AUTHOR CONTRIBUTIONS

Y.B. conceived the study and prepared the figures. Y.B. and M.N. wrote the manuscript. M.N, S.S. and Y.B. designed the experiments. M.N. performed the following experiments: hiTSC and hiPSC reprogramming, immunostaining, immunohistochemistry, directed differentiation, CRISPR/Cas9-mediated SOX2 KO HFFs, PI FACS analysis, organoid culture. A.R. analyzed the RNA-seq, RRBS data, ATAC-seq and ChIP-seq data. V.Z. cloned the various transcription factors and initiated the reprogramming experiments toward hiTSCs. S.S. prepared the RNA libraries, performed immunohistochemistry, ELISA, qPCR for C19MC miRNAs and subcutaneously injected cells into NOD/SCID mice. R.L., N.D. and D.O. ran qPCRs and validated SOX2 KO. K.M. injected hiTSCs. M.J. prepared the DKO cells and run reprogramming experiments. O.S. and H.C. prepared the RRBS libraries and analyzed the RRBS data. M.R. performed part of the organoid experiments and qPCR. H.Y. generated the episomal-derived hiTSCs. A.K grow hiTSCs and collected samples for further analysis. S.EL. and R.E. derived the hbdTSC lines. M.NP., D GW., and S.Y. provided with critical trophoblast knowledge and helped in designing the differentiation experiments.

## DATA AND SOFTWARE AVAILABILITY

All RNA-seq, RRBS, ATAC-seq and ChIP-seq data were deposited in the Gene Expression Omnibus database (GEO) under accession number GSE182017 (https://www.ncbi.nlm.nih.gov/geo/query/acc.cgi?acc=GSE182017). The authors declare no competing financial interests. Correspondence and requests for materials should be addressed to Y.B.

## METHODS

### Derivation of human trophoblast stem cells from human blastocysts and human fibroblasts from skin biopsy and cell lines

The establishment and use of hbdTSC lines or hESC lines from PGD-derived embryos was performed in compliance with the protocols approved by the Ethic Committee of Shaare Zedek Medical Center (IRB 87/07). Embryo donations were carried out under the strict regulation of the National ethic committee (Israel health ministry) and NIH and ISSCR guidelines. In order to generate human blastocyst-derived TSC (hbdTSC) control lines, human blastocysts were plated on Mitomycin C-treated mouse embryonic fibroblast (MEF) feeder cells and cultured in human TSC medium as previously described^1^. Following blastocyst outgrowth, the cells were trypsinized and transferred into new Mitomycin C-treated MEF feeder plates. The cells were passaged several times, until stable proliferative hbdTSCs emerged. PCS201 human primary fibroblasts were purchased from ATCC (PCS-201-012). GM2 female adult fibroblasts (GM25432) were a gift from Dr. Oren Ram (Hebrew University of Jerusalem). These cells were derived by Coriell INSTITUTE (https://catalog.coriell.org/) in compliance with their regulations. The derivation of KEN human fibroblasts from foreskin biopsy was performed in compliance with protocols approved by the Ethic Committee of Shaare Zedek Medical Center (IRB 88/11). hESCs was provided by Dr. Rachel Eiges and the breast cancer cell line MDA-MB-231 (HTB-26) was purchased from ATCC.

### Molecular Cloning and hiTSC and hiPSC reprogramming

All dox-inducible factors were generated by cloning the open reading frame of each factor into the pMINI vector (NEB) and then restricted with EcoRI or MfeI and inserted into the FUW-TetO expression vector. A lentiviral vector dox-dependent system was utilized for the transient expression of transcription factors. For infection, replication-incompetent lentiviruses containing the various reprogramming factors (GOKM for hiTSC reprogramming and OKSM for hiPSC reprogramming) were packaged with a lentiviral packaging mix (7.5μg psPAX2 and 2.5μg pDGM.2) in HEK 293T cells and collected 48, 60, 72 and 84 hours after transfection. The supernatants were filtered through a 0.45μm filter, supplemented with 8μg/ml of polybrene, and then used to infect fibroblasts which were previously infected with lentiviral vectors encoding puromycin resistance and M2rtTA, and subsequently selected with 2 µg/ml puromycin for 3-5 days. Twelve hours following the fourth infection, medium was replaced with basic reprogramming medium (BRM) consisting of fresh DMEM containing 10%FBS, 1% L-glutamine and 1% penicillin-streptomycin.

For hiTSC reprogramming, six hours after medium replacement 2μg/ml doxycycline (dox) was added to the medium. The reprogramming medium was changed every other day. After 14 days in BRM with dox, the medium was replaced with 50% BRM and 50% hTSC medium ^1^ with dox, followed by 7 days in hTSC medium with dox. Then, dox was removed and colonies were allowed to stabilize for 7-10 days. Plates were then screened for primary hiTSC colonies. Each colony was isolated by manually picking up colonies with a pipette, trypsinized with TrypLE (Gibco) and plated in a separate well on feeder cells in hTSC medium. The cells were passaged several times until stable proliferative hiTSC colonies emerged.

For hiPSC reprogramming, six hours after medium replacement 2μg/ml doxycycline (dox) was added to the medium. The reprogramming medium was changed every other day. After 7 days in BRM, the medium was replaced with 50% BRM and 50% human embryonic stem cell (hES) medium comprised of Knockout DMEM containing 15% KnockOut serum replacement, 0.1 mM 2-mercaptoethanol, 1% L-glutamine, 1% non-essential amino acids and 1% Penicillin-Streptomycin, with dox, followed by 7 additional days in hES medium with dox. Then, dox was removed and colonies were allowed to stabilize for 7-10 days in hES medium supplemented with freshly added 4ng/ml bFGF. Each colony was isolated by manually cutting colonies into small chunks with a Pasteur pipette and manually transferring each with a pipette to a separate well on feeder cells in hES medium supplemented with 4ng/ml bFGF freshly added to each well.

### Generation of non-integrating episomal-derived hiTSCs

Episomal vectors: pCLXE-hOCT4-shp53, pCXWB-EBNA1, pCLXE-hL-MYC/LIN28A vectors used for this study were originally described in (Okita et al. 2011) (Okita et al. 2012) and were obtained from Addgene (Cat 27078, Cat 37624, and Cat 27077, respectively). pCXLE-hGATA3 and pCXLE-hKLF4 vectors were generated by subcloning the open reading frame of either hGATA3 or hKLF4 into pCXLE vector using EcoRI enzyme. For electroporation, fibroblasts were trypsinized and 3.5×10^5^ cells were resuspended in 100 µL of R buffer (Life Technologies, Carlsbad, CA, USA). Subsequently, 1µg/100µL of each of the episomal plasmids (pCXLE-hGATA3, pCXLE-hKLF4, pCXLE-hUL) with 1.5µg/100 µL of pCXLE-hOCT4-shp53 and 0.5µg/100 µL of pCXWB-EBNA1 episomal plasmids were added to the cell suspension. Electroporation was carried out with the NEON transfection system according to the manufacturer’s instructions (Life Technologies; 1,650V, 20ms, 1 pulse). Subsequently, cells were seeded onto 6-well plates containing basic reprogramming medium (BRM) consisting of fresh DMEM containing 10%FBS, 1% L-glutamine, and 1% penicillin-streptomycin. For hiTSC reprogramming, the fibroblasts were cultured in BRM for 10 days, and then the medium was exchanged to hTSC medium (Okae et al., 2018) for another 14 days. Plates were then screened for primary hiTSC colonies. Each colony was isolated by manually picking up colonies with a pipette, trypsinized with TrypLE (Gibco), and plated in a separate well on feeder cells in hTSC medium. The cells were passaged several times until stable proliferative hiTSC colonies emerged.

### Quantitative PCR (qPCR) for mRNA expression and analysis of genomic integration of transgenes

For analysis of mRNA expression using qPCR, total RNA was isolated using the Macherey-Nagel kit (Ornat). 500–1000ng of total RNA was reverse transcribed using iScript cDNA Synthesis kit (Bio-Rad). Quantitative PCR analysis was performed in duplicates using 1/100 of the reverse transcription reaction in a StepOnePlus (Applied Biosystems) with SYBR green Fast qPCR Mix (Applied Biosystems). Specific primers were designed for the different genes (see Table 1). All quantitative real-time PCR experiments were normalized to the expression of *GAPDH* and presented as a mean ± standard deviation of two duplicate runs.

**Table 1.**
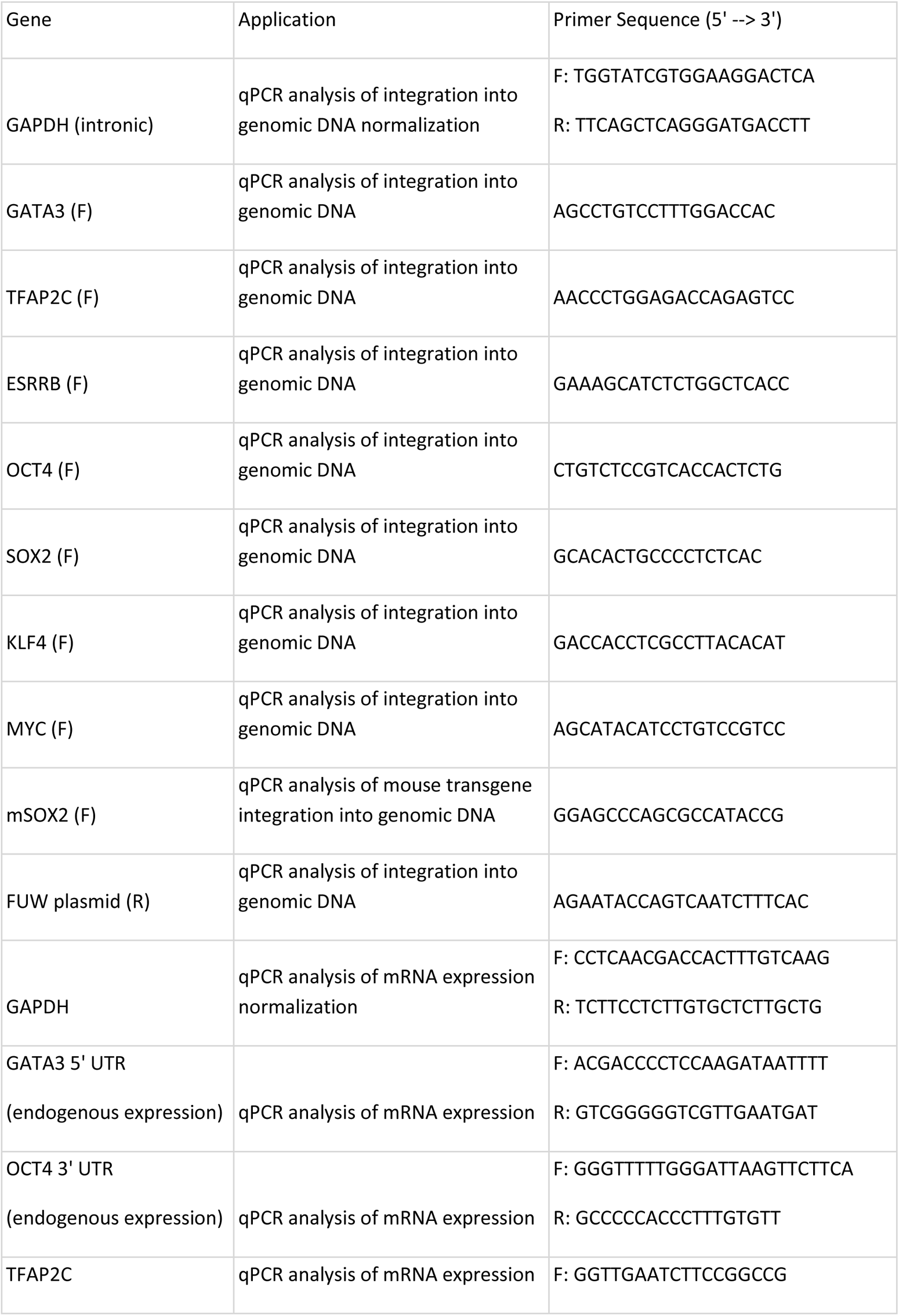
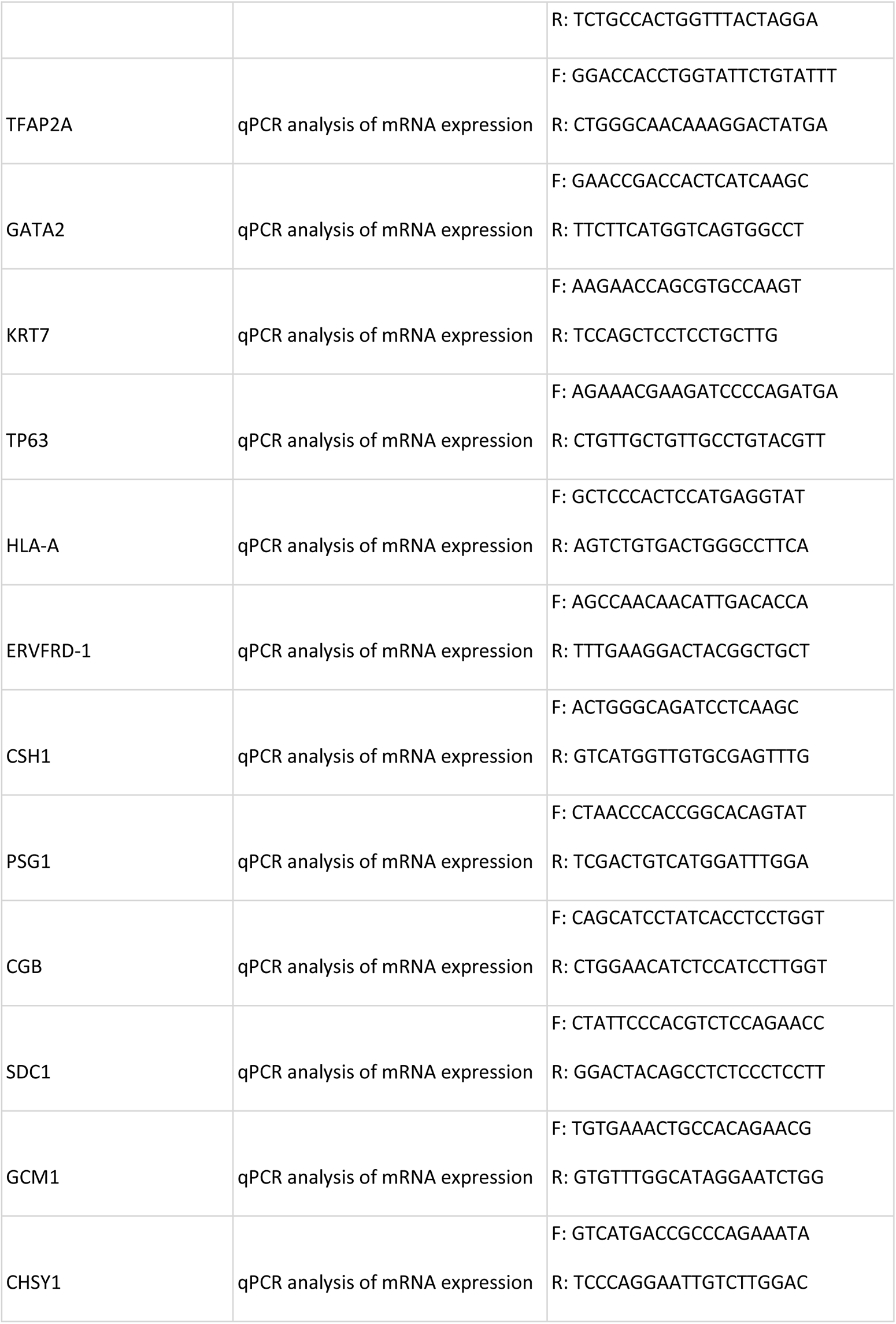
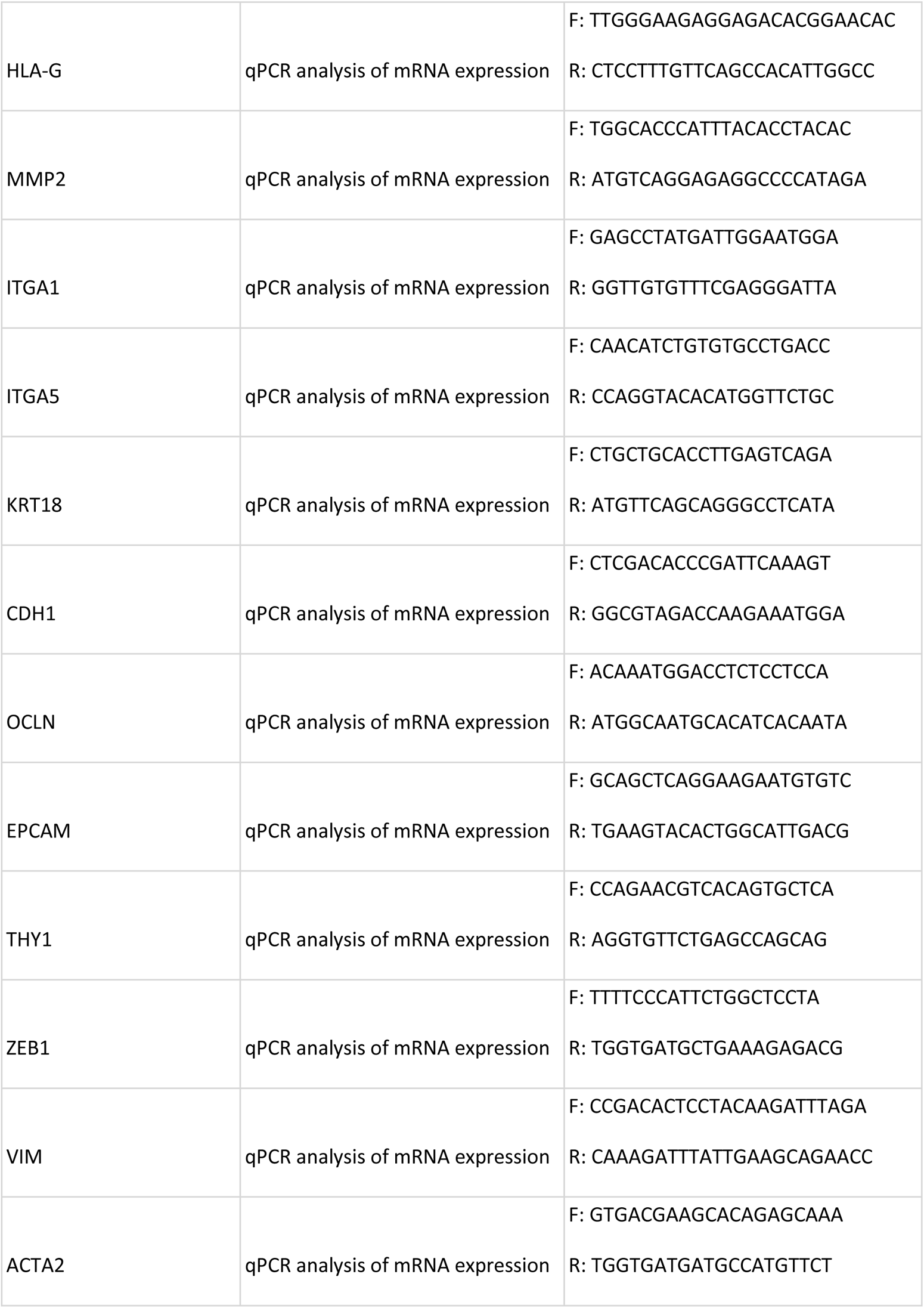
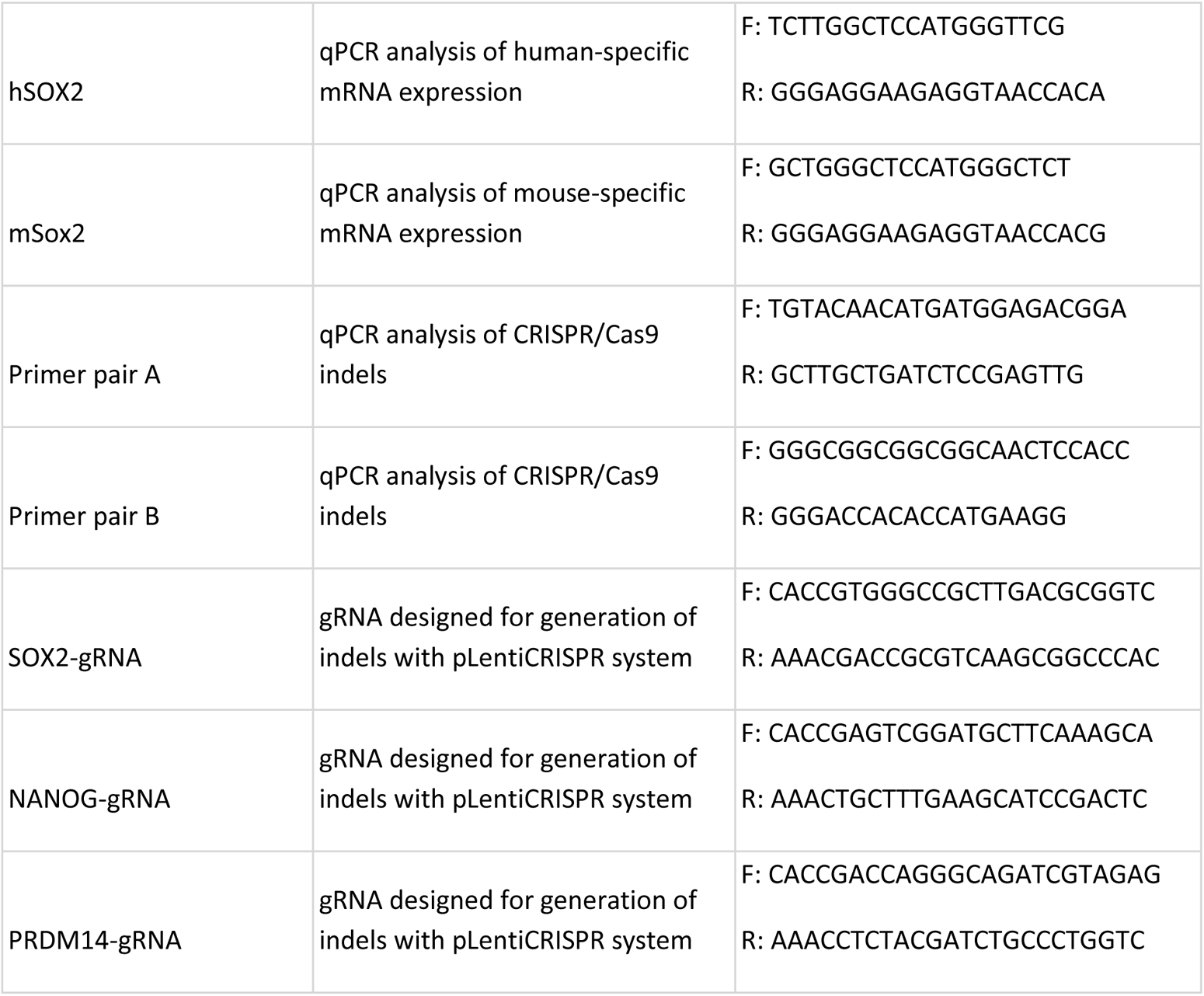
primer list

For analysis of integration of transgenes into genomic DNA using qPCR, genomic DNA was isolated by incubating trypsinized cell pellets in lysis buffer consisting of 100mM Tris pH8, 5mM EDTA, 0.2% SDS and 200mM NaCl overnight with 400μg/ml proteinase K (Axxora) at 37°C for one hour followed by incubation at 55°C for one hour. Then, genomic DNA was precipitated with isopropanol, washed with 70% ethanol and resuspended in ultra-pure water (BI). Forward primers for the end of the last exon of cloned genes were used in conjunction with reverse primers for the FUW-tetO vector at the region immediately downstream of the cloned gene (see Table 1). Results were normalized to an intronic region of the *GAPDH* gene and presented as a mean ± standard deviation of two duplicate runs.

### qRT-PCR of miRNA

Cells of four hiTSC colonies, hbdTSC, hESCs and breast cancer cell line MDA-MB-231 were lysed and total RNA was isolated using the Macherey-Nagel kit (Ornat). To quantify C19MC miRNAs, we adapted a previously published method ^51^. Briefly, RNA (10 ng) in 15 ml reaction mixture was converted into cDNA using RT primers (50 nM) that were complementary to each miRNA with a TaqMan MicroRNA Reverse Transcription Kit (Life Technologies #4366596). Primers were adapted from (Lee et al., 2016). The cDNAs were quantified by qRT-PCR with qPCRBIO Fast qPCR SyGreen (Tamar # PB20.16). hsa-miR-103a was used for normalization of the results ^23^.

### Immunostaining of PFA-fixated cells and flow cytometry

Cells were fixed in 4% paraformaldehyde (in PBS) for 20 minutes, then rinsed 3 times with PBS and blocked for 1 hour with PBS containing 0.1% triton X-100 and 5%FBS. The cells were incubated overnight with primary antibodies (1:200-1:500 dilution) in 4C. The antibodies are: anti-KRT7 (Abcam, ab215855), anti-GATA3 (Abcam, ab106625), anti-GATA2 (Abcam, ab173817), anti-TFAP2C (Santa Cruz Biotechnologies, sc-12762), anti-KRT18 (Santa Cruz Biotechnologies, sc-51582), anti-CDH1 (Santa Cruz Biotechnologies, sc-7870), anti-VIM (Cell Signaling Technology, #5741), anti-SDC1 (Abcam, ab128936), anti-CSH1 (Abcam, ab15554), anti-HLA-G (Abcam, ab52455), anti-ITGA5 (Abcam, ab150361), anti-SOX2 (Abcam, ab97959), anti-OCT4 (Abcam, ab19857), anti-TRA-1-60 (Abcam, ab16288) diluted in PBS containing 0.1% triton X-100 and 1% FBS. The next day, the cells were washed 3 times and incubated for 1 hour with relevant (Alexa) secondary antibody in PBS containing 0.1% triton X-100 and 1%FBS (1:500 dilution). DAPI was added 10 minutes before end of incubation. Negative control included incubation with secondary antibody without primary.

For flow cytometry analysis of HLA class I and TRA-1-60 expression, cells were trypsinized and blocked for ten minutes in incubation buffer containing 0.5% bovine serum albumin (BSA) (Sigma Aldrich) in PBS. Then, cells were centrifuged and resuspended in incubation buffer with anti-HLA class I (1:300, Abcam, ab22432) or anti-TRA-1-60 (1:300, Abcam, ab16288) for 1 hour. Cells were then washed with incubation buffer and incubated for 30 minutes with relevant (Alexa) secondary antibody, after which cells were washed, resuspended in incubation buffer, filtered through mesh and analyzed and/or sorted. HLA class I stained cells were analyzed by FACS (Beckman Coulter). Results were analyzed using Kaluza Software. TRA-1-60 stained cells were sorted using FACSAria III (BD Biosciences).

### RNA and RRBS library preparation and sequencing and karyotype analysis

For RNAseq, total RNA was isolated using the Qiagen RNeasy kit. mRNA libraries were prepared using the SENSE mRNA-seq library prep kit V2 (Lexogen), and pooled libraries were sequenced on an Illumina NextSeq 500 platform to generate 75-bp single-end reads.

For RRBS, DNA was isolated from samples and incubated in lysis buffer (25mM Tris-HCl at pH8, 2mM EDTA, 0.2%SDS, 200mM NaCl) supplemented with 300μg/mL proteinase K (Roche) followed by phenol:chloroform extraction and ethanol precipitation. hiTSC colonies and hbdTSC colonies were passaged twice on matrigel in order to eliminate the presence of MEF feeder cells. RRBS libraries were prepared as previously described ^52^. Samples were run on HiSeq 2500 (Illumina) using 100bp paired-end sequencing.

Karyotype analysis was performed on identical isolated DNA samples using Affymetrix CytoScan 750K array.

### Chromatin immunoprecipitation (ChIP)

Chromatin immunoprecipitation (ChIP) assay was performed as previously described ^53^. Briefly, cells from two biological replicates per line were fixed for 10min at RT with a final formaldehyde concentration of 0.8%. Formaldehyde was quenched with glycine at a final concentration of 125mM. The cells were then lysed with lysis buffer (100mM Tris-HCl, 300mM NaCl, 2% Triton X-100, 0.2%v sodium deoxycholate and 10mM Cacl2) supplemented with EDTA-free protease inhibitor (Roche, 11873580001) for 20min on ice. The chromatin was digested by adding MNase (Thermo Scientific, 88216) for 20 min at 37°C and MNase was inactivated by adding 20mM EGTA. The fragmented chromatin was added to pre-bounded Dynabeads (A and G mix, Invitrogen, 10004D/ 10002D) using H3K4me2 antibody (Millipore, 07-030) at 2 μg per reaction. Samples were then washed twice with RIPA buffer, twice with RIPA high salt buffer (NaCl 360mM), twice with LiCl wash buffer (10mM Tris-Hcl, 250mM LiCl, 0.5% DOC, 1mM EDTA, 0.5% IGEPAL) and twice with 10mM Tris-HCl pH= 8. DNA was purified by adding RNase A (Thermo Scientific, EN0531) and incubated for 30 min at 37°C and then with Proteinase K (Invitrogen, 25530049) for 2h. The DNA was eluted by adding 2X concentrated elution buffer (10mM Tris-HCl, 300mM NaCl, 1% SDS, 2mM EDTA) and reverse crosslinked overnight at 65°C. The DNA was then extracted using AMPure XP beads (Beckman Coulter Genomics, A63881). ChIP sample libraries were prepared according to Illumina Genomic DNA protocol.

### ATAC libraries and Sequencing

ATAC-Seq library preparation was performed as previously described ^54^. Briefly, 50,000 cells per replicate (two biological replicates per line) were incubated with 0.1% NP-40 to isolate nuclei. Nuclei were then transposed for 30 min at 37C with adaptor-loaded Nextera Tn5 (Illumina, Fc-121-1030). Transposed fragments were directly PCR amplified and Sequenced on an Illumina NextSeq 500 platform to generate 2 x 36-bp paired-end reads.

### Differentiation of hiTSCs

For directed differentiation into ST, approximately 10^5^ cells were seeded on Matrigel coated 12-well plates at a concentration of 1:30 in ambient oxygen conditions in a medium consisting of DMEM/F12 supplemented with 0.1mM 2-mercaptoethanol, 0.5% Penicillin-Streptomycin, 0.3% BSA, 1% ITS supplement, 2.5 μM Y27632, 2 μM forskolin, and 4% KSR, as described ^1^. Cells were collected at day 2 and 6 for analysis of mRNA expression using qPCR as described above. Cells were also seeded on 12-well plates at a density of approximately 10^5^ cells per plate, cultured similarly and fixated in 4% PFA for immunostaining as described above.

For directed differentiation into EVT, approximately 4×10^5^ cells were seeded on Matrigel coated 12-well plates at a concentration of 1:100 in ambient oxygen conditions in a medium consisting of DMEM/F12 supplemented with 0.1mM 2-mercaptoethanol, 0.5% Penicillin-Streptomycin, 0.3% BSA, 1% ITS supplement, 100 ng/ml NRG1, 7.5μM A83-01, 2.5μM Y27632, and 4% KnockOut Serum Replacement, as described by ^1^. Matrigel was added to a final concentration of 2%. At day 3, the medium was replaced with the EVT medium without NRG1, and Matrigel was added to a final concentration of 0.5%. Medium was replaced every other day, and cells were collected at day 7 and 14. Cells were also fixated in 4% PFA for immunostaining as described above.

### Formation of trophoblast organoids with bdTSCs and hiTSCs

Similar to as described in ^42^, bdTSCs and hiTSCs were suspended in trophoblast organoid medium (TOM) consisting of DMEM/F12, 10mM HEPES, 1× B27, 1× N2, 1mM L-glutamine, 100ng/mL R-spondin, 1μM A83-01, 100ng/mL recombinant human epidermal growth factor (rhEGF), 50ng/mL recombinant murine hepatocyte growth factor (rmHGF), 2.5μM prostaglandin E2, 3μM CHIR99021, and 100ng/mL Noggin. Growth factor-reduced Matrigel (GFR-M) was added to reach a final concentration of 60%. Solution (40μL) containing 10^4^-10^5^ bdTSCs/hiTSCs was placed in the center of 24-well plates. After 2 min at 37°C, the plates were turned upside down to ensure equal spreading of the cells in the solidifying GFR-M-forming domes. After 15min, the plates were turned again and the domes were carefully overlaid with 500 μL prewarmed TOM. Cells were cultured in 5% oxygen for 10-19 days and then subject to immunostaining.

### Immunostaining of bdTSC and hiTSC trophoblast organoids

Organoid-containing Matrigel domes were fixated in 4% PFA overnight. Then, domes were washed with PBS for 15 minutes twice. Domes were submerged in blocking solution containing 3% bovine serum albumin (BSA), 5% fetal bovine serum (FBS), 0.1% Triton X-100 in PBS, at 4C overnight. Then, tissues were incubated with primary antibodies including anti-Ki67 (1:200 Abcam, ab15580), anti-KRT7 (1:200, Abcam, ab215855), anti-CSH1 (1:200, Abcam, ab15554) and anti-TFAP2C (1:100, Santa Cruz Biotechnologies, sc-12762) diluted in PBS containing 1% BSA and 0.1% Triton X-100, on a rocking plate at 4C for two nights. Plates were moved to room temperature and continued rocking for at least 2 additional hours before washing in PBS containing 0.1% Triton X-100 overnight, with at least 5 changes of buffer. Following this, domes were incubated in secondary antibody solution containing relevant (Alexa) secondary antibody (1:200) diluted in 1% BSA and 0.1% Triton X-100 on a rocking plate at 4C overnight. Then, domes were washed again with PBS containing 0.1% Triton X-100 overnight, with at least 5 changes of buffer. Domes were then incubated with DAPI for 1 hour and stored in PBS in 4C until imaging. Imaging was performed using spinning disk confocal microscopy with Nikon Eclipse Ti2 CSU-W1 Yokogawa confocal scanning unit, Andor Zyla sCMOS camera and Nikon Plan Apo VC 20X NA 0.75 lens. Maximal intensity projection images were created using NIS-Elements microscope imaging software.

### Engraftment of hiTSCs into NOD/SCID mice and immunohistochemistry (IHC)

For each lesion, approximately 4×10^6^ were trypsinized with TrypLE, washed twice in PBS, resuspended in 150μl of a 1:2 mixture of Matrigel and PBS and subcutaneously injected into NOD/SCID mice.

The joint ethics committee (IACUC) of the Hebrew University and Hadassah Medical Center approved the study protocol (IACUC# MD-18-15628-3) for animal welfare. The Hebrew University is an AAALAC international accredited institute.

Lesions were collected nine days after injection, dissected, fixed in 4% paraformaldehyde overnight, embedded in paraffin, sectioned and mounted onto slides. Some slides were stained with H&E, while others were subject to IHC staining.

For IHC, slides were deparaffinized in xylene and rehydrated in a decreasing ethanol gradient. Antigen retrieval was performed in a sodium citrate buffer and slides were heated for 3 minutes at 110-120C. After a short incubation in 3% hydrogen peroxide, sections were incubated overnight in CAS-block (Invitrogen) with primary antibodies anti-KRT7 (1:1000, Abcam, ab215855), HLA-G (1:100, Abcam, ab52455) and anti-CSH1 (1:100, Abcam, ab15554). Then, sections were incubated with appropriate HRP-conjugated secondary antibody (Vector Laboratories) for 30 minutes and immunohistochemistry was performed using DAB peroxidase substrate kit (Vector Laboratories). Slides were lightly counterstained with hematoxylin.

### hCG detection in the blood of hiTSC-injected mice

For testing presence of hCG in NOD/SCID mice, approximately 500μl of mouse serum was collected in Eppendorf tubes by collecting whole blood and centrifuging at 1500G for 10 minutes at 4°C to separate serum. Serum samples were stored at −80°C until processing.

Quantification of human chorionic gonadotropin (hCG) hormone levels were determined by ELISA kit by Alpco (Almog diagnostic #25-HCGHU-E01), following the manufacturer’s protocol.

### ELF5 bisulfite sequencing

ELF5 bisulfite sequencing was performed as previously published ^23^. DNA from each sample was treated with bisulfite using the EpiTect Bisulfite Kit (Qiagen #59110), according to the manufacturer’s protocol. 10% of the resulting DNA was used for the amplification of the 432 to 3 bp region upstream of the ELF5 start site via nested PCR. Amplicons were directly sequenced using MiSeq 2×150 bp paired-end run.

### Generation of SOX2 and NANOG/PRDM14 KO hiTSCs

gRNAs designed to target the first exon of SOX2, NANOG and PRDM14 (see Table 1) were cloned into pLentiCRISPR V2 vectors with puro (SOX2, NANOG) or hygromycin B (PRDM14) resistance (Addgene plasmids #52961 and #98291 respectively) using BsmBI restriction enzyme, as described in ^47^. These were used to infect HFFs (KEN), which were previously infected with lentiviral vectors encoding for GFP and M2rtTA. Next, cells were selected with 2 µg/ml puromycin for 3-5 days (SOX2-KO), or underwent double selection with both 2 µg/ml puromycin for 3-5 days and 200 ug/ml hygromycin B for 5-7 days (NANOG/PRDM14-DKO). Identical fibroblasts infected with pLentiCRISPR V2 vectors without gRNA and similarly selected were used as controls. Cells were then subjected to reprogramming to hiTSCs or hiPSCs using GOKM or OSKM factors, as described above. For analysis of the presence of genomic indels, as well as analysis of mouse vs human SOX2 integration and expression, colonies were passaged at least twice on Matrigel 1:30 without the presence of MEF feeder cells. For assessment of indel events in hiTSCs resulting from the SOX2-KO HFF reprogramming, genomic DNA was isolated as described above and analyzed using PCR and running products on agarose gel, or qPCR with specific primer pairs (see Figure 7B and Table 1). For assessment of indel events in hiTSCs resulting from the NANOG/PRDM14-DKO HFF reprogramming, colonies were subjected to Sanger sequencing at the NANOG and PRDM14 gene loci (HAI Laboratories) and visualized using Sequencher software.

### RNA-Seq analysis

For analysis of RNA-seq data, raw reads (FastQ files) were quality-trimmed using Trim Galore (v 0.6.5, default parameters, http://www.bioinformatics.babraham.ac.uk/projects/trim_galore) and aligned to the human genome GRCh38 using HISAT2 (v 2.1.0, http://daehwankimlab.github.io/hisat2/). Data were quantitated at mRNA level using htseq-counts (v 0.13.5). Genes with a sum of counts less than 10 over all samples were filtered out, retaining 230408 genes. Differential expression analysis was performed using the DESeq2 package (v 1.28.1, ^55^). Gene Ontology Enrichment Analysis using R package enrichR (v3.0, https://maayanlab.cloud/Enrichr/) for top 1000 genes that are up-regulated in all GOKM hiTSCs colonies compared to fibroblasts. Network analysis for top 2000 up-regulated genes between hiTSCs and fibroblasts. Protein-protein interaction network was analyzed with STRING (http://www.string-db.Org). The MCODE plugin tool (http://baderlab.org/Software/MCODE) in Cytoscape was used to extract densely connected sub-networks. For each subnetwork we used iRegulon plugin tool (http://iregulon.aertslab.org/) in Cytoscape to analyze systematically the composition of the gene promoters in transcription factor binding sites.

### DNA Methylation analysis

For the analysis of RRBS data, raw reads (FastQ files) were quality-trimmed using Trim Galore (v 0.6.5, default parameters) and aligned to the human genome GRCh38 using BSMAP (v 2.9 ^56^). The methylation ratio of CpGs with sequencing depth of at least 10 reads were computed based on 100bp tiles. Differentially methylated regions (DMR) table obtained from Methylkit (v 1.14.2, DOI: <u>10.18129/B9.bioc.methylKit</u>) processing of the BAM files yielded by BSMAP alignment. Each table represents the following parameters: Chromosome (chr), start and end coordinates of the methylated region (start, end), strand location (strand), probability value “pvalue”, adjusted p-value “qvalue”, differential methylation score “meth.diff”. Only regions with a meth.diff score over 50 or under −50 and a q-value under 1E-5 where considered as differentially methylated (hyper- or hypo-methylated respectively).

For the analysis of direct amplification and sequencing of the *ELF5 promoter a* FATSTA file that contains ~20 kbp from the human genome GRCh38 upstream of ELF5 (chr11:34496606-34517332) was constructed. An index for the ELF5 FASTA file was constructed using the function “bismark_genome_preparation” from the Bismark (v 0.22.3, ^57^). Alignment was then performed using bismark --bowtie2 ^58^ and methylation. Finally, methylation information was extracted using the function “bismark_methylation_extractor”.

### ATAC-Seq and ChIP-Seq analysis

For analysis of the accessibility and activity data, raw reads (FastQ files) were quality-trimmed using Trim Galore (v0.6.5, default parameters) and aligned to the human genome GRCh38 using burrows-wheeler aligner (BWA v 0.7.17, ^59^). Peaks were called with MACS2 (v 2.2.7.1 ^60^). Differential peak analysis was performed using the DESeq2 package (v 1.28.1, DOI: <u>10.18129/B9.bioc.DESeq2</u>). Peaks were defined specifically for each group of cells and the R package (Venn v 1.9) was used to construct all Venn diagrams. Peaks that are specific to OSKM D3 and OGKM D3 were differentially identified with FDR< 0.05. Intersection of these peaks with hbdTSCs and hESCs allowed for the identification of specific set of peaks that were investigated for possible regulation of nearby genes. GREAT ^61^ was used to associate these peaks with genes within 250 kb. Extracted genes were then intersected with a specific set of genes that were differentially expressed in hbdTSCs and not in fibroblasts or hESCs, or set of genes that were differentially expressed in hESCs and not in fibroblasts or hbdTSCs. Cytoscape plugin iRegulon was used to identify possible regulators of those genes (FDR <0.05 and NES > 2). Transcription factors were added to the network iteratively based on whether they are overexpressed from the beginning of the reprogramming process and then if a specific transcription factor was detected at day 3 of the reprogramming process. Finally, if we find a significant number of genes that are not attached to the other nodes in the network we assume that another set of transcription factors might be involved in the regulation of these genes later in the process.

## SUPPLEMENTARY DATA

**Figure S1.**
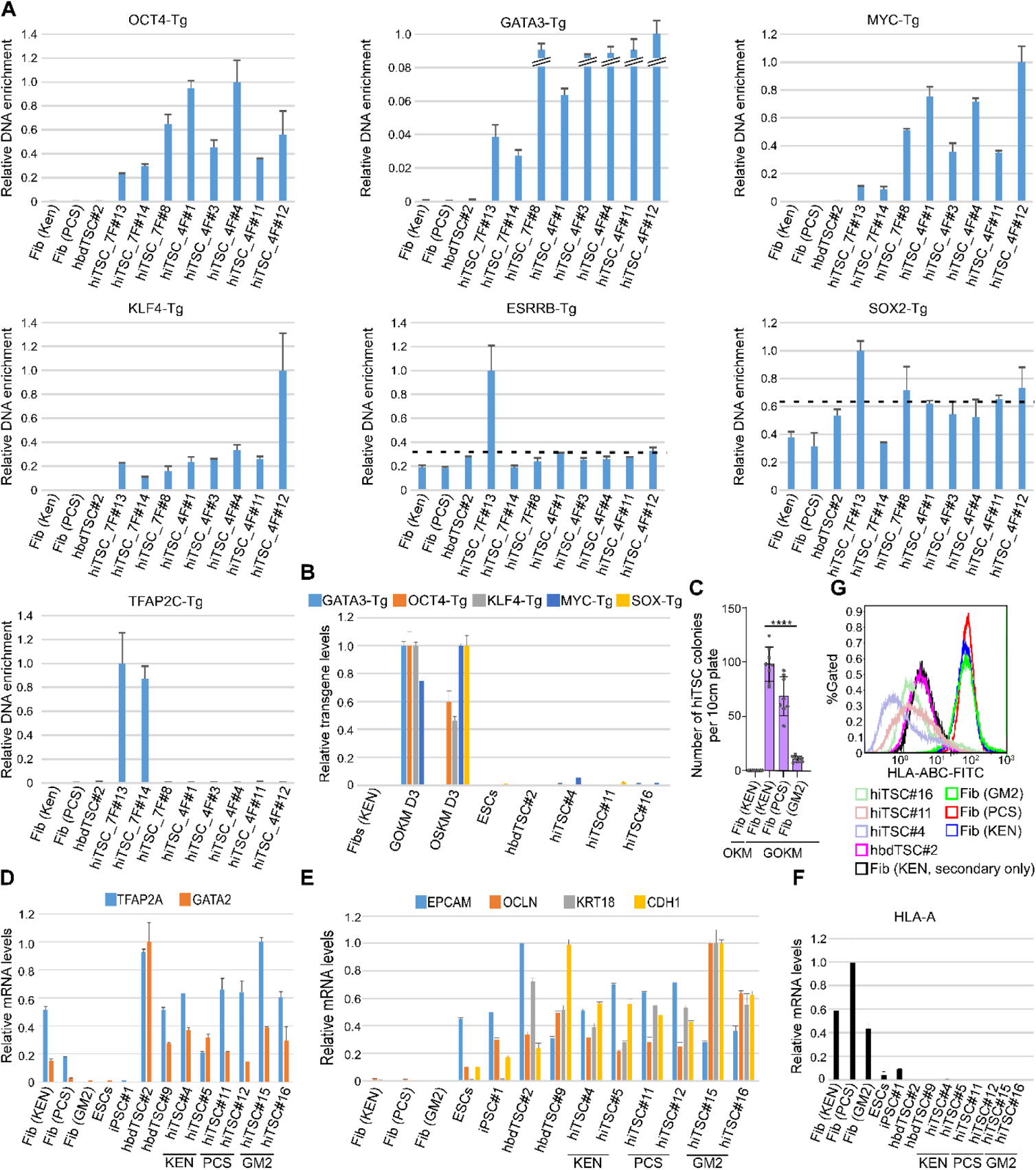
Fibroblasts reprogrammed to hiTSCs with forced expression of GOKM undergo MET and express hTSC markers. **(A)** GATA3, OCT4, KLF4 and MYC transgens are present in all hiTSC colonies. qPCR analysis of the indicated transgenes in hiTSC colonies and negative controls HFFs (KEN), HFFs (PSC) and hbdTSC#2. Transgene integration was assessed by forward primers designed for the last exon of the transgene with reverse primers matching the sequence of the FUW-tetO plasmid (See Table 1). The highest sample for each transgene was set to 1. Results were normalized to an intronic region of the *GAPDH* gene and are shown as fold change. Error bars indicate standard deviation between two duplicates. Dotted lines mark the negative threshold of the various transgenes based on uninfected cells (i.e. fibroblasts or hbdTSCs). **(B)** hiTSC colonies exhibit minor to no transgene leakiness. qPCR analysis of mRNA levels of the indicated transgenes in various hiTSC colonies and in GOKM or OSKM infected fibroblasts following 3 days of dox exposure. Uninfected fibroblasts and hbdTSCs were used as negative controls for transgene expression. The highest sample for each transgene was set to 1. Results were normalized to the mRNA levels of the housekeeping control gene *GAPDH* and are shown as fold change. Error bars indicate standard deviation between two duplicates. hiTSC colonies were derived from 3 independent reprogramming experiments. **(C)** hiTSC reprogramming efficiency varies between different primary fibroblast lines. Graph showing an average number of hiTSC colonies generated by either OKM or GOKM in the indicated fibroblast lines. Error bars indicate standard deviation between 8 replicates. ****p-value< 0.001 as calculated by GraphPad Prism using Student’s t-test. **(D)** qPCR analysis of mRNA levels of trophoblast markers *GATA2* and *TFAP2A* in the indicated hbdTSC clones, hiTSC colonies, fibroblast lines and pluripotent stem cell (PSC) lines. Note that *TFAP2A* and to some extent *GATA2* are expressed in fibroblasts. The highest sample for each gene was set to 1. Results were normalized to the mRNA levels of the housekeeping control gene *GAPDH* and are shown as fold change. Error bars indicate standard deviation between two duplicates. **(E)** qPCR analysis of mRNA levels of epithelial markers *EPCAM, OCLN, KRT18, and CDH1* in the indicated hbdTSC clones, hiTSC colonies, fibroblast lines and PSC lines. The highest sample for each gene was set to 1. Results were normalized to the mRNA levels of the housekeeping control gene *GAPDH* and are shown as fold change. Error bars indicate standard deviation between two duplicates in a typical experiment. **(F)** qPCR analysis of mRNA levels of HLA class I gene *HLA-A* in the indicated samples. The highest sample was set to 1. Results were normalized to the mRNA levels of the housekeeping control gene *GAPDH* and are shown as fold change. Error bars indicate standard deviation between two duplicates. **(G)** Flow cytometry histogram of classical HLA class I protein expression (HLA-A/B/C) in the indicated hiTSC colonies, hbdTSC#2 and fibroblast lines using the well-characterized W6/32 antibody. Fibroblasts (KEN) stained with only secondary antibody used as negative control.

**Figure S2.**
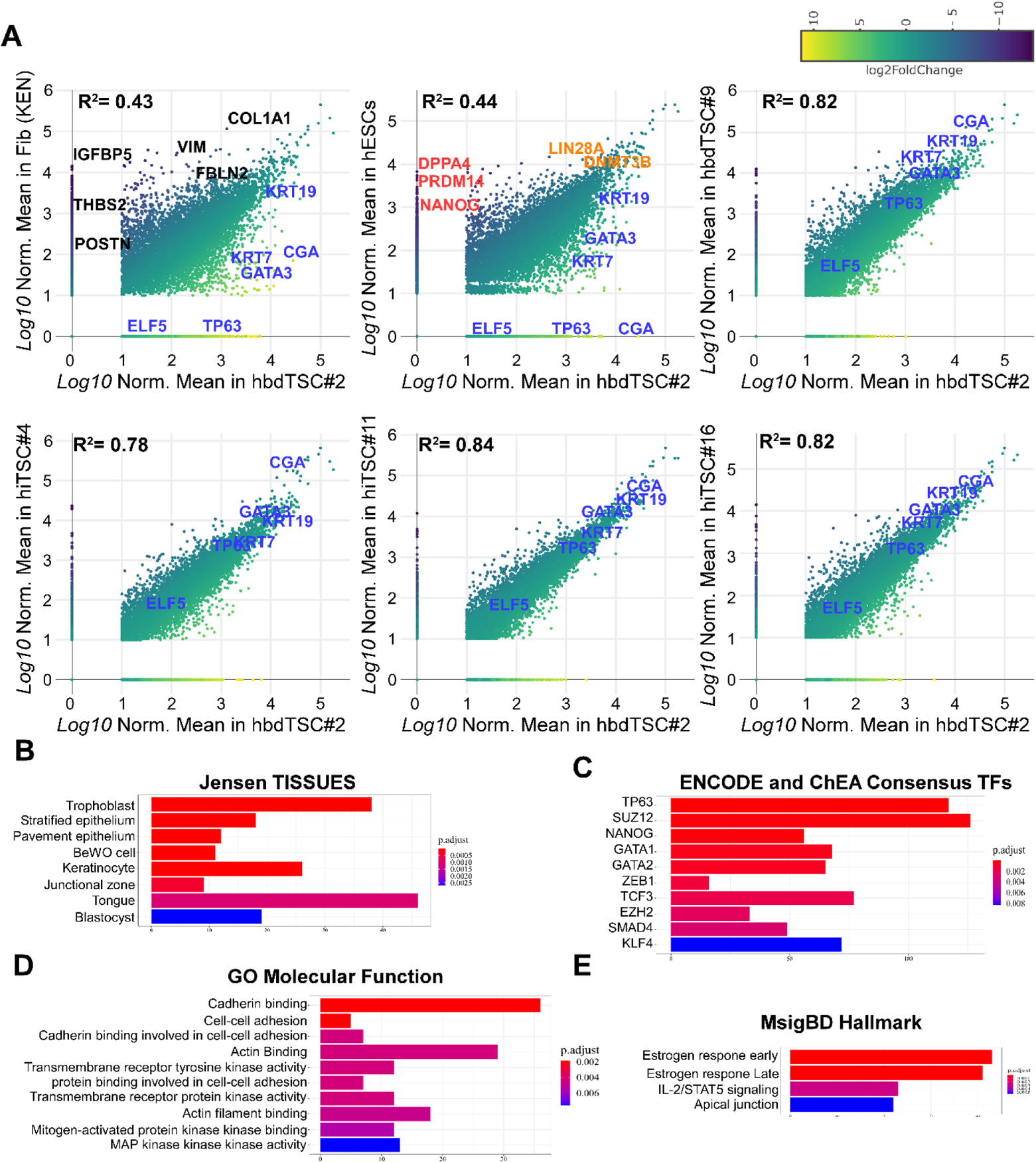
RNA-seq analysis indicates that hiTSCs have a transcriptome enriched for gene ontology terms related to placental development. **(A)** Scatter plots displaying pairwise correlations of gene expression levels for hbdTSC#2 vs KEN fibroblasts, hbdTSC#2 vs hESCs, hbdTSC#2 vs hbdTSC#9, hbdTSC#2 vs hiTSC#4, hbdTSC#2 vs hiTSC#11, and hbdTSC#2 vs hiTSC#16. Two RNA-seq replicates were generated for differential gene expression analyses (LFC> 2, p.adj < 1E-3, Log_10_ Normalized Mean counts > 1). Each dot represents an individual gene, while color scale indicates LFC (Log_2_ fold change). Genes that are marked in blue or black indicate genes that are specific to hTSCs and fibroblasts, respectively. Genes that are marked in red indicate genes that are specific to hESCs, while genes that are shared between hbdTSCs and hESCs are marked in orange. R^2^ value was calculated for each pairwise comparison, demonstrating a high degree of similarity between hiTSCs and hbdTSCs. **(B-E)** Bar graph showing the highest enriched GO terms for top 1000 most differentially expressed genes between hiTSCs and fibroblasts under the Jensen Tissues, ENCODE and ChEA Consensus TFs, GO Molecular Function and MsigBD Hallmark categories. Jensen Tissues database showed trophoblast as the highest enriched term (p.adj < 1E-12), while TP63, NANOG and GATA2 are among the top 10 enriched transcription factors (p.adj< 0.0005).

**Figure S3.**
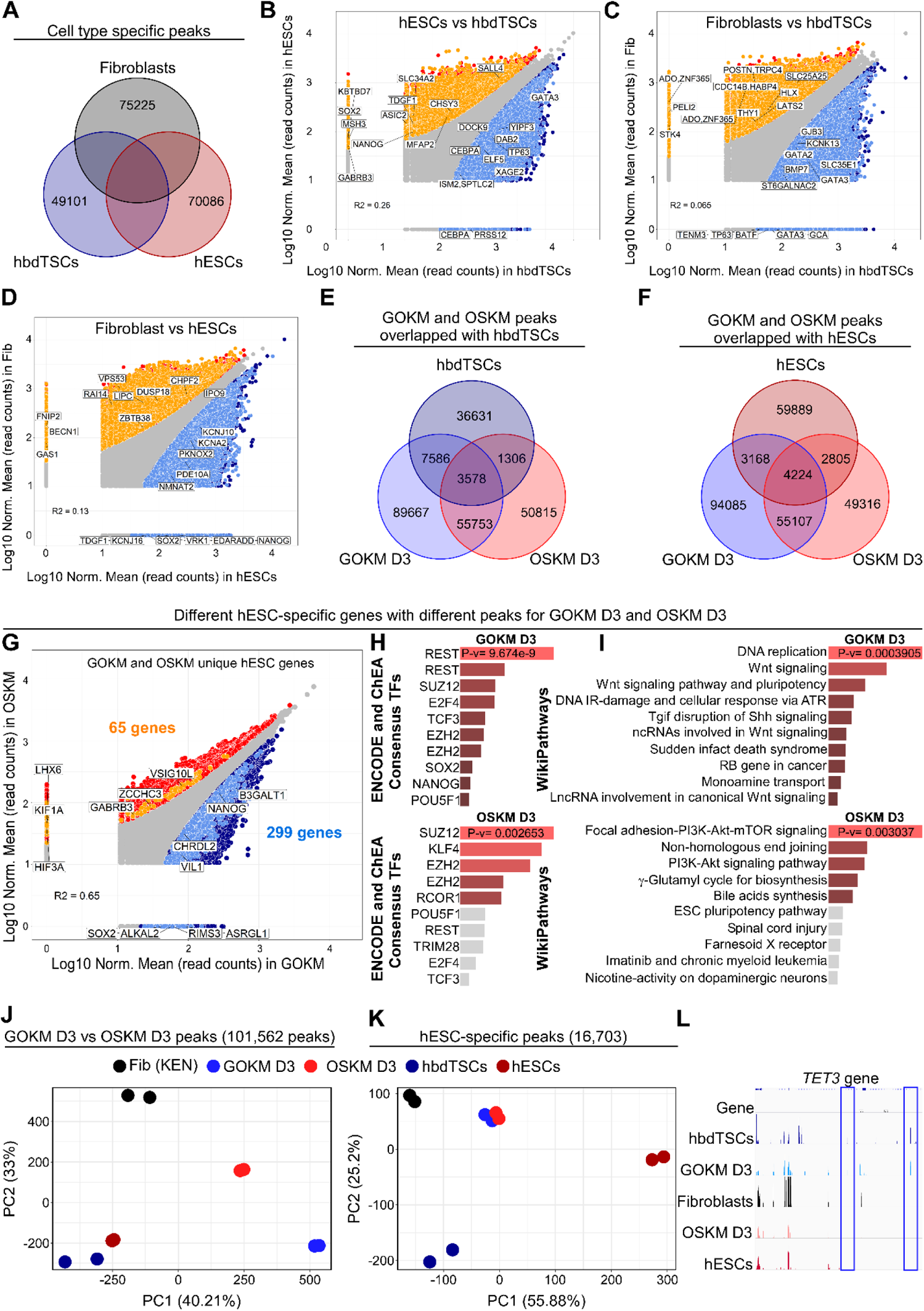
ATAC-seq analysis on GOKM and OSKM-transduced cells following 3 days of transgene induction. **(A)** Venn diagram of ATAC-seq showing peaks that are specific to fibroblasts, hbdTSCs, and hESCs (FDR< 0.01). **(B-D)** Scatter plots displaying pairwise correlations and differentially accessible peaks between hESCs vs hbdTSCs (B), fibroblasts vs hbdTSCs (C), and fibroblasts vs hESCs (D) with FDR< 0.01. Peaks that are specific to differential peaks along the horizontal axis or the vertical axis were labeled by dark blue and red, respectively. Peaks associated with genes that are specific to the samples along the horizontal axes were labeled with light blue, while peaks associated with genes that are specific to samples along the vertical axes were labeled with light orange. **(E-F)** Venn diagrams of ATAC-seq showing the overlap between peaks that are specific to hbdTSCs (E) or hESCs (F) with differentially accessibility between GOKM D3, OSKM D3 with FDR< 0.05. **(G)** A scatter plot of differentially accessible peaks between GOKM D3, OSKM D3 with FDR< 0.05. Peaks that are specific to GOKM D3 or OSKM D3 are labeled by dark blue and red, respectively. Peaks associated with genes specific to the GOKM reprogramming and are specific to the hESC state were labeled with light blue, while peaks associated with genes specific to the OSKM reprogramming and are specific to the hESC state are labeled with orange. 299 genes are found to overlap between hESC-specific genes and peaks that are associated to these genes specifically in GOKM D3, while 65 genes are found to overlap between hESC-specific genes and peaks that are associated with these genes in OSKM D3. **(H-I)** GO terms for 299 and 65 genes in (G) under the categories “ENCODE and ChEA Consensus TFs” and “WikiPathways” respectively. **(J)** PCA analysis of 5 different groups across 101,562 accessibility regions that are differential between OSKM D3 and GOKM D3. A significant change in chromatin accessibility is observed in GOKM D3 compared to OSKM D3. **(K)** PCA analysis of 5 different groups across 16,703 accessibility regions that are specific to hESCs. GOKM D3 and OSKM D3 clustered together indicating that the major differences between the two reprogramming models are specific to the hbdTSCs specific peaks and not due to hESCs peaks. **(L)** Integrative Genomics Viewer of snapshots of ATAC-seq signal accessible regions at the *TET3 gene locus*. Blue rectangle marks peaks that are shared between GOKM D3 and hbdTSCs but not with OSKM D3, parental fibroblasts or hESCs.

**Figure S4.**
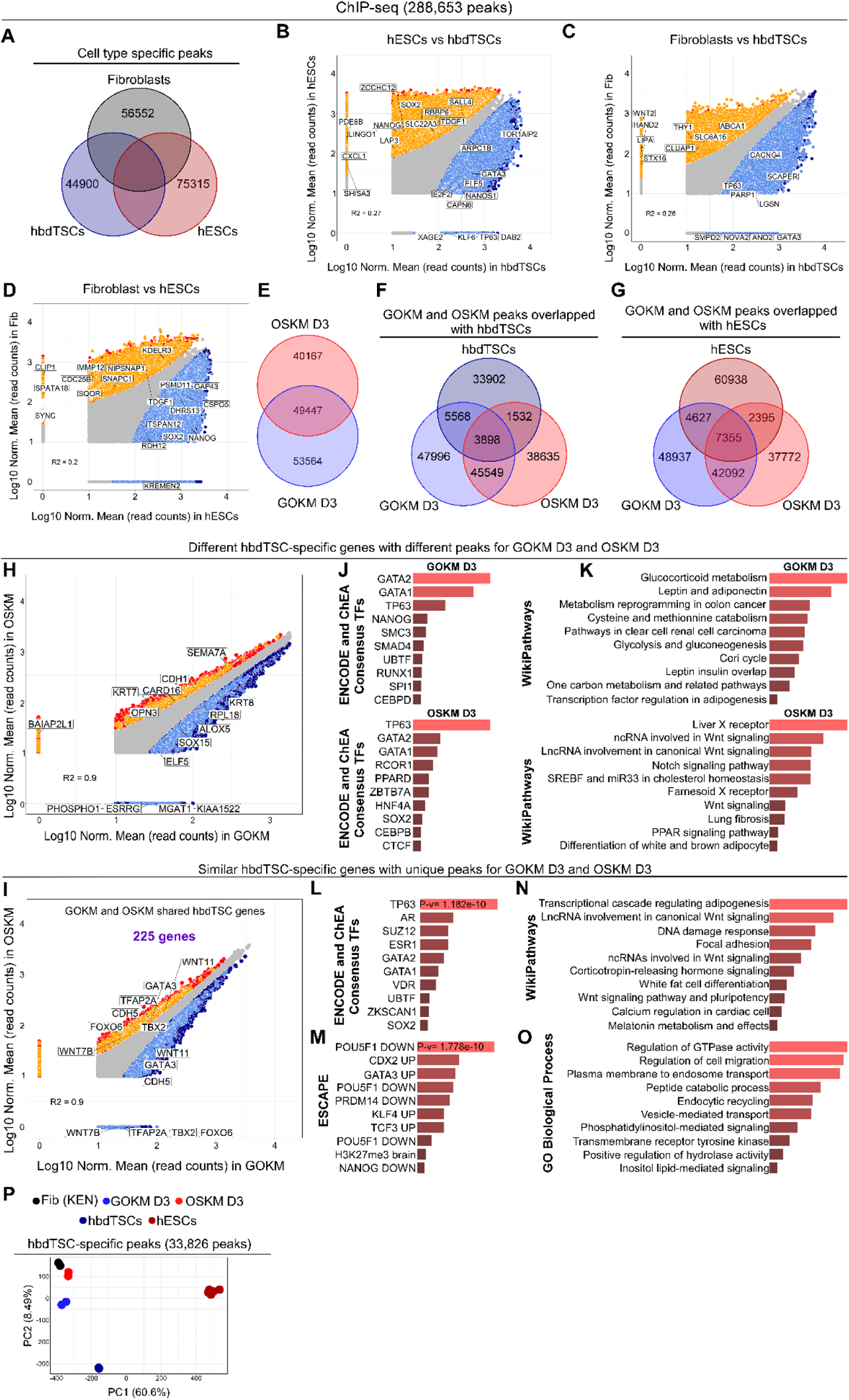
ChIP-seq analysis for H3K4me2 on GOKM and OSKM-transduced cells following 3 days of transgene induction. **(A)** Venn diagram of ChIP-seq for H3K4me2 showing peaks that are specific to fibroblasts, hbdTSCs, and hESCs (FDR< 0.01). **(B-D)** Scatter plots displaying pairwise correlations and differentially accessible peaks between hESCs vs hbdTSCs (B), fibroblasts vs hbdTSCs (C), and fibroblasts vs hESCs (D) with FDR< 0.01. Peaks that are specific to differential peaks along the horizontal axis or the vertical axis were labeled by dark blue and red, respectively. Peaks associated with genes that are specific to the samples along the horizontal axes were labeled with light blue, while peaks associated with genes that are specific to samples along the vertical axes were labeled with light orange. **(E-G)** Venn diagrams of ChIP-seq showing the overlap between peaks that are specific to hbdTSCs (F) or hESCs (G) with differentially activity between GOKM D3, OSKM D3 with FDR< 0.05. **(H)** A scatter plot of differentially H3K4me2 deposition between GOKM D3 and OSKM D3 with FDR< 0.05. Peaks that are specific to GOKM D3 or OSKM D3 are labeled by dark blue and red, respectively. Peaks associated with genes specific to the GOKM reprogramming and are specific to the hESC state are labeled with light blue, while peaks associated with genes specific to the OSKM reprogramming and are specific to the hESC state are labeled with orange. 183 genes are found to overlap between hbdTSC-specific genes and peaks that are associated with these genes in GOKM D3, while 161 genes are found to overlap between hbdTSC-specific genes and peaks that are associated with these genes in OSKM D3. **(J-K)** GO terms for the 183 and 161 genes in (H) under the categories “ENCODE and ChEA Consensus TFs” and “WikiPathways”. **(I)** A scatter plot of differentially H3K4me2 deposition peaks between GOKM D3 and OSKM D3 with FDR< 0.05. Peaks that are specific to GOKM or OSKM are labeled by dark blue and red respectively. Peaks associated with genes that are shared between the hbdTSC state and in day 3 of the GOKM and OSKM reprogramming are labeled with light blue, or with orange, respectively. 225 genes are shared between the two reprogramming systems with different H3K4me2 deposition. **(L-O)** GO terms for the 225 genes in (I) under the categories “ENCODE and ChEA Consensus TFs” (L), “ESCAPE” (M), “WikiPathways” (N), and GO biological process (O). **(P)** PCA analysis of 5 different groups across 33,826 H3K4me2 deposited regions that are differential between GOKM D3 and OSKM D3 (FDR< 0.05). A clear separation between the two groups is observed, putting GOKM D3 in a closer distance toward the ‘hbdTSCs’ sample.

**Figure S5.**
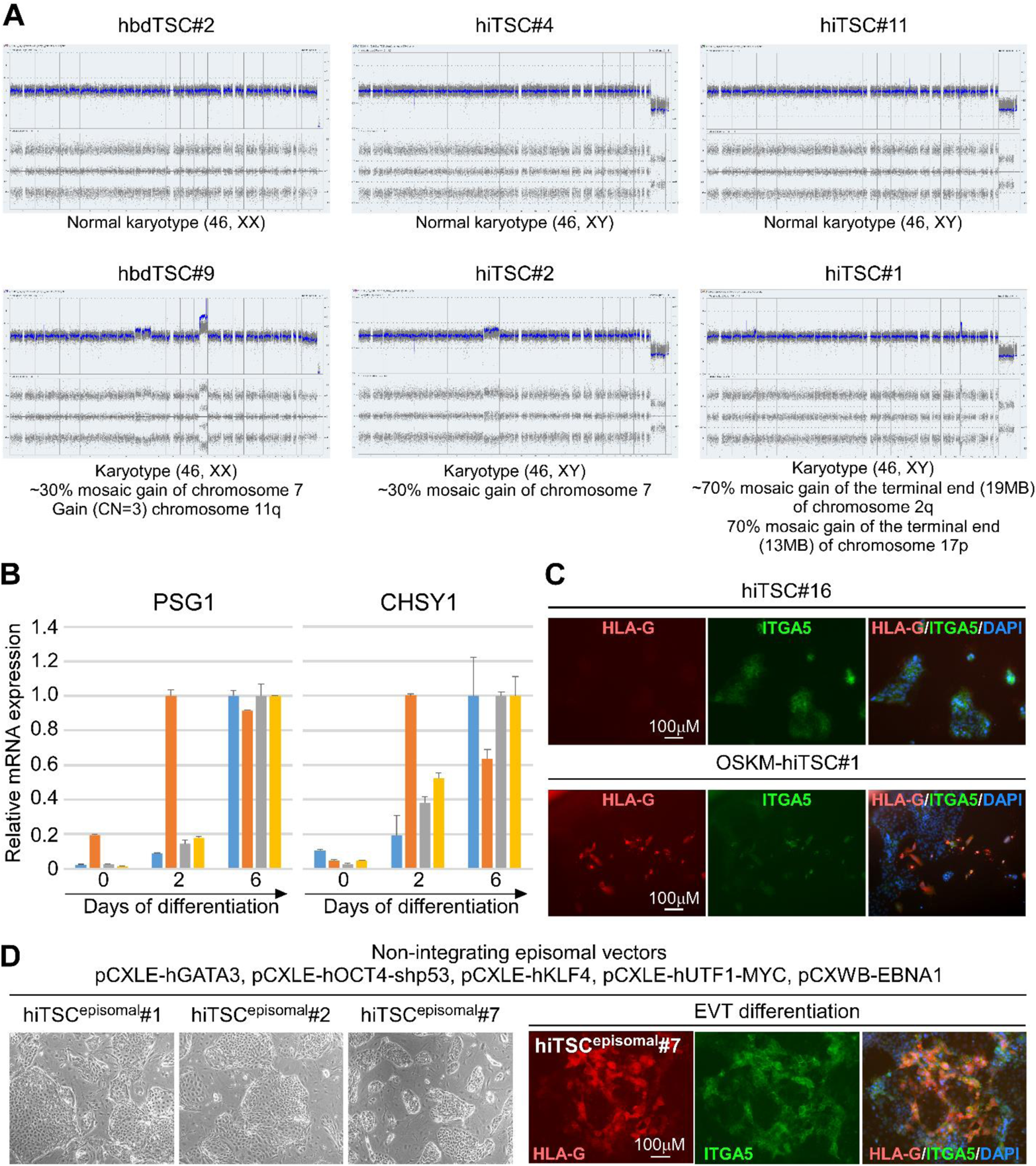
hiTSCs can maintain normal karyotype and differentiate into ST and EVT-like cells. Both hbdTSC#2 and hbdTSC#9 were isolated from PGD embryos (see Methods). Geneic examination of the two lines reveals that hbdTSC#2 is heterozygous for RB mutation and hbdTSC#9 is heterozygous for RB and Marfan mutations **(A)** Plots displaying the karyotype of hbdTSC and hiTSC lines. Two hbdTSC lines, hbdTSC#2 and hbdTSC#9, and four hiTSC clones, hiTSC#1, hiTSC#2, hiTSC#4 and hiTSC#11, were subjected to karyotyping analysis using Affymetrix CytoScan 750K array. 50% of hbdTSC lines and 50% of hiTSC lines displayed an intact chromosomal karyotype. The other 50% of the colonies exhibited few aberrations in a fraction of the cells. The specific aberrations and the relevant affected fraction of the cells are specified below each plot. **(B)** qPCR analysis of relative mRNA levels of ST marker genes *PSG1* and *CHSY1* at days 0, 2 and 6 in ST differentiation protocol. The highest sample for each gene in each colony was set to 1. Results were normalized to the mRNA levels of the housekeeping control gene *GAPDH* and are shown as fold change. Error bars indicate standard deviation between two duplicates. **(C)** Immunofluorescent staining for the EVT-specific markers HLA-G and ITGA5 and DAPI nuclear staining in PFA-fixated hiTSC#16 and OSKM-hiTSC#1 following 14 days of EVT differentiation. **(D, left)** Brightfield images of isolated hiTSC clones derived using the indicated episomes, delivered through electroporation. **(D, right)** Immunofluorescent staining for the EVT-specific markers HLA-G and ITGA5 and DAPI nuclear staining in PFA-fixated hiTSC^episomal^#7 following 14 days of EVT differentiation.

**Figure S6.**
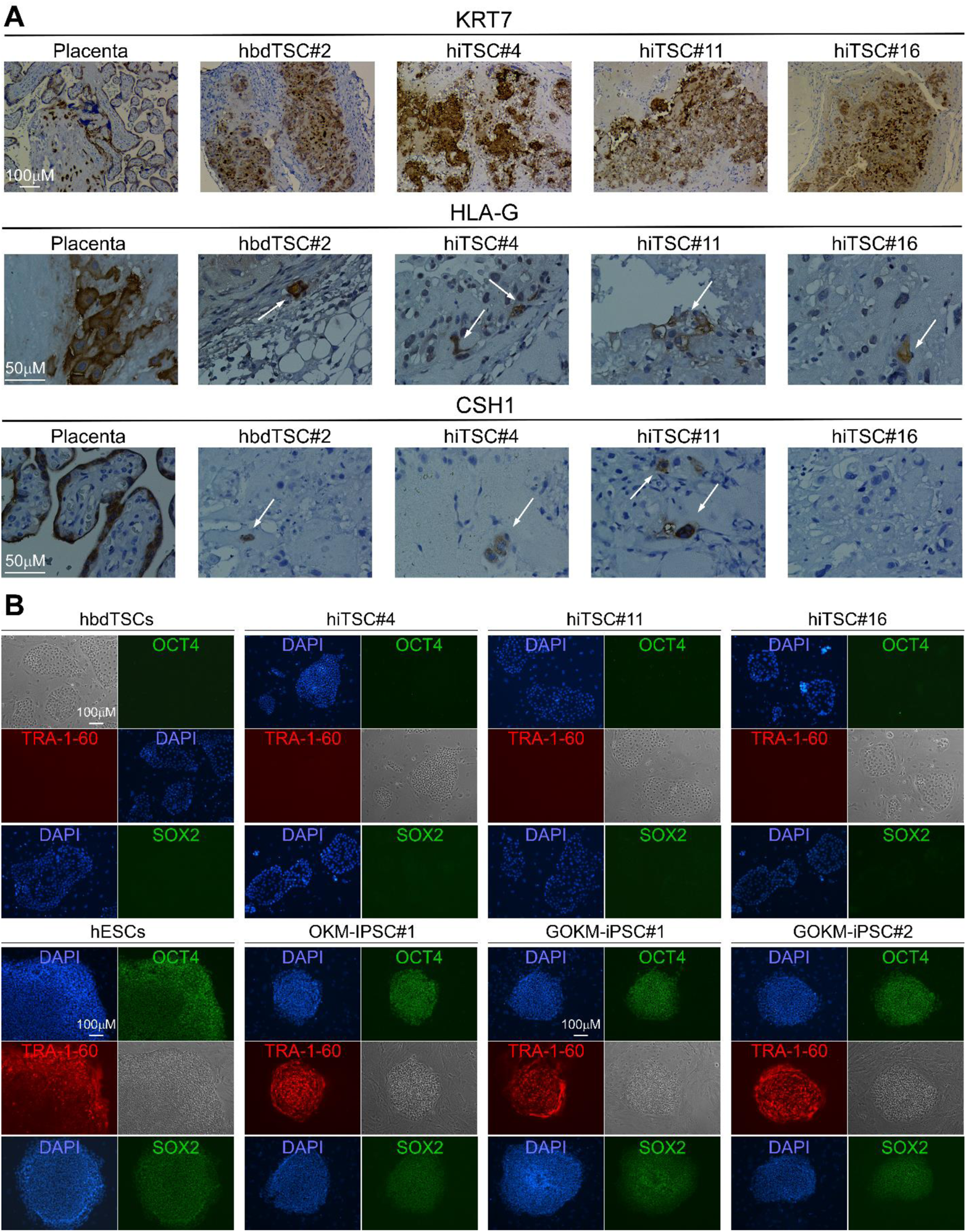
hiTSCs engrafted into NOD-SCID mice form trophoblastic lesions and hiTSCs are negative to pluripotency-specific markers. For lesion formation approximately 4×10^6^ hbdTSCs or hiTSCs were subcutaneously injected into NOD-SCID mice. Lesions were collected nine days after injection and analyzed by immunohistochemistry for specific markers. **(A)** Immunohistochemically stained sections of human placenta and trophoblastic lesions extracted from SCID-NOD mice showing strong KRT7 staining (top) and scattered staining for the EVT marker HLA-G (middle) and ST marker CSH1 (bottom). White arrows point to positive staining for the indicated markers **(B)** hiTSCs are negative to pluripotent-specific markers. Fluorescent images displaying high expression levels of OCT4, SOX2 and TRA-1-60 in pluripotent stem cells (hESCs, GOKM-hiPSCs and OKM-hiPSCs, bottom panel) and none (OCT4 and TRA-1-60) to very low expression (SOX2) in hiTSCs (upper panel).

**Figure S7.**
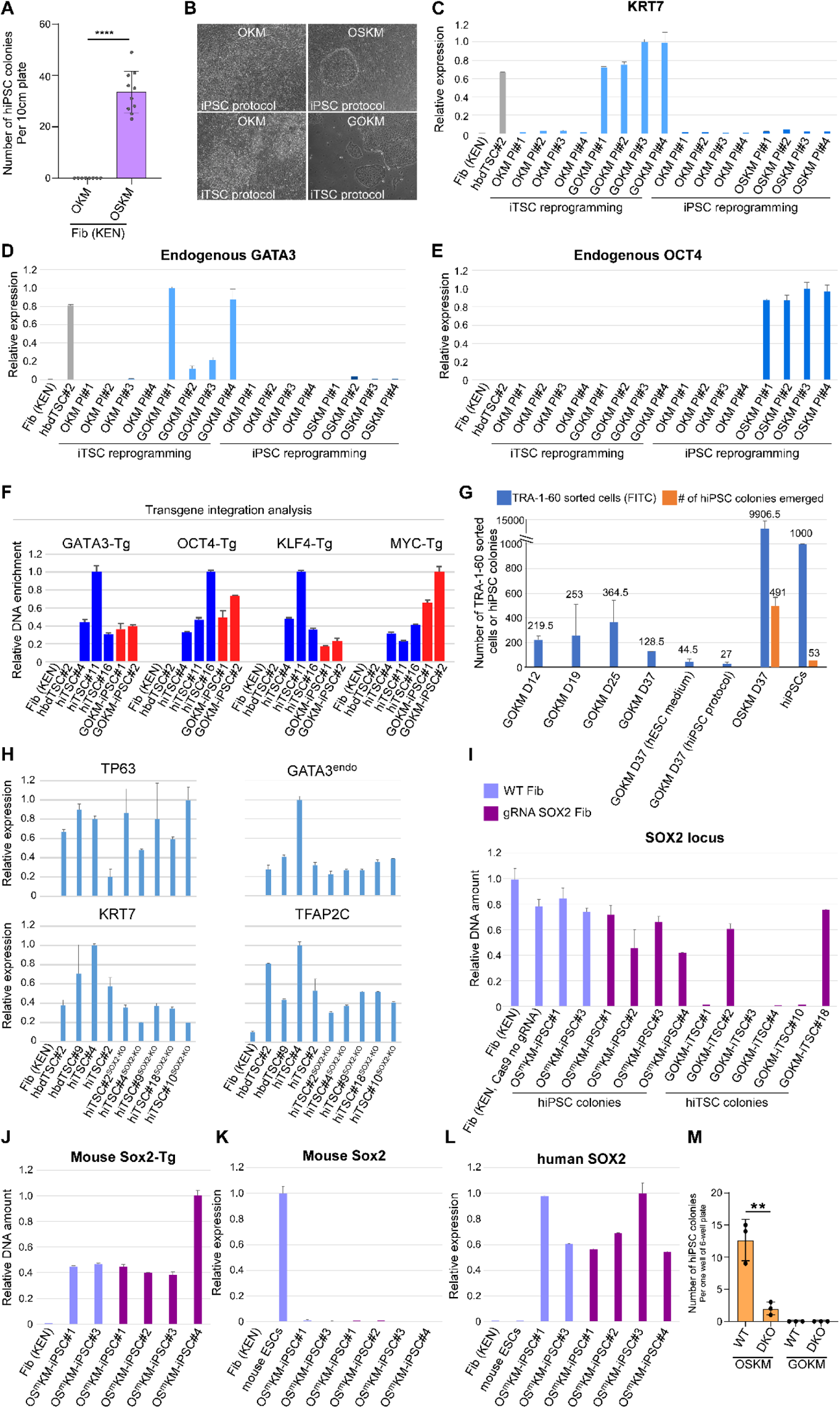
OKM are incapable of generating hiTSCs and GOKM do not acquire pluripotency during hiTSC formation. Fibroblasts were transduced with OKM, GOKM or OSKM and reprogramming efficiency in generating iPSCs or hiTSCs was examined **(A)** Graph showing an average number of hiPSC colonies generated by either OKM or OSKM. Error bars indicate standard deviation between 8-10 replicates. ****p-value< 0.001 as calculated by GraphPad Prism using Student’s t-test. Graph depicting the average number of hiTSC colonies generated by OKM and GOKM is located in Figure S1C. Note, following multiple attempts we were able to derive two OKM and two GOKM-hiPSC colonies (See Figure S6B). **(B, top)** Bright field images of representative areas in plates which were reprogrammed with OKM or OSKM using hiPSC reprogramming protocol (left) and following colony isolation (right). **(B, bottom)** Bright field images of representative areas in plates which were reprogrammed with OKM or GOKM using hiTSC reprogramming protocol (left) and following colony isolation (right). **(C-E)** qPCR analysis of mRNA levels of trophoblast markers *KRT7* (C) and endogenous UTR expression of *GATA3* (D) and the pluripotency marker endogenous UTR expression of *OCT4* (E) in plates (4 each) transduced with OKM, GOKM or OSKM following hiPSC or hiTSC reprogramming protocols as specified in the graphs. The highest sample for each gene was set to 1. Results were normalized to the mRNA levels of the housekeeping control gene *GAPDH* and are shown as fold change. Error bars indicate standard deviation between two duplicates. **(F)** qPCR analysis for the relative DNA enrichment of GOKM transgene integration in the indicated samples. Note, the two GOKM-iPSC colonies contain low levels of KLF4-Tg and high levels of MYC-Tg compared to GOKM-hiTSC colonies. Transgene integration was assessed using forward primers designed for the last exon of the transgene with reverse primers matching the sequence of the FUW-TetO plasmid (See Table 1). The highest sample for each transgene was set to 1. Results were normalized to unaffected genomic region of the *GAPDH* gene and are shown as fold change. Error bars indicate standard deviation between two duplicates. **(G)** Graph displaying the average number of TRA-1-60-positive cells that were sorted during GOKM-mediated hiTSC or hiPSC reprogramming protocols at indicated time points, and the number of hiPSC colonies that emerged in each plate. OSKM-mediated hiPSC reprogramming at day 37 and stable hiPSCs were used as positive control. Error bars indicate standard deviation between two biological replicates. **(H)** qPCR analysis of mRNA levels of TSC-specific markers *TP63*, *GATA3^endo^*, *KRT7* and *TFAP2C* in Fibroblast (KEN), two hbdTSC clones, two hiTSC clones and five SOX2 KO hiTSC clones. The highest sample for each gene was set to 1. Results were normalized to the mRNA levels of the housekeeping control gene *GAPDH* and are shown as fold change. Error bars indicate standard deviation between two duplicates. **(I)** qPCR analysis for the relative DNA enrichment of the human *SOX2* genomic locus in six OKM+ mouse Sox2 (OS^m^KM)-derived hiPSC colonies and six GOKM-derived hiTSC colonies in either WT fibroblasts (light purple) or fibroblasts transduced with lentiviral vector encoding for puromycin resistance and constitutively active Cas9 together with a gRNA against the SOX2 coding region (dark purple). WT fibroblast (KEN) and fibroblasts transduced with lentiviral vector encoding for puromycin resistance and constitutively active Cas9, but without gRNA, were used as negative controls. Note that while 4 out of 6 GOKM-derived hiTSC clones displayed homozygous deletions for the human *SOX2* genomic locus, all the examined OS^m^KM-derived hiPSC clones exhibited at least one intact SOX2 allele (using primer pair B, see Figure 7 and Table 1). The highest sample for each gene was set to 1. Results were normalized to unaffected genomic region of the *GAPDH* gene and are shown as fold change. Error bars indicate standard deviation between two duplicates. **(J-L)** qPCR analysis for the relative DNA enrichment of the mouse *Sox2 transgene* (J), mRNA levels of mouse *Sox2* gene (K) and mRNA levels of human *SOX2* gene (L) in the six OS^m^KM-derived hiPSC clones (from I) and fibroblast controls. Note that all six OSKM-derived hiPSC clones display mouse *Sox2* transgene integration in their genome without leaky expression and high levels of endogenous human SOX2 gene expression. The highest sample for each gene was set to 1. Results were normalized to unaffected genomic region of the *GAPDH* gene and are shown as fold change. Error bars indicate standard deviation between two duplicates. **(M)** Graph displaying the number of hiPSC colonies that were generated following reprogramming with pluripotency protocol using OSKM or GOKM factors in WT or double knockout (DKO, for NANOG and PRDM14) fibroblasts. Error bars indicate standard deviation between 3 replicates. **p-value < 0.01 as calculated by GraphPad Prism using Student’s t-test.

## Notes

### Competing Interest Statement

The authors have declared no competing interest.

